# Oscillatory cortical forces promote three dimensional cell intercalations that shape the mandibular arch

**DOI:** 10.1101/309120

**Authors:** Hirotaka Tao, Min Zhu, Kimberly Lau, Owen K.W. Whitley, Mohammad Samani, Xiao Xiao, Xiao Xiao Chen, Noah A. Hahn, Weifan Liu, Megan Valencia, Min Wu, Xian Wang, Kelli D. Fenelon, Clarissa C. Pasiliao, Di Hu, Jinchun Wu, Shoshana Spring, James Ferguson, Edith P. Karuna, R. Mark Henkelman, Alexander Dunn, Huaxiong Huang, Hsin-Yi Henry Ho, Radhika Atit, Sidhartha Goyal, Yu Sun, Sevan Hopyan

## Abstract

Multiple vertebrate embryonic structures such as organ primordia are composed of a volume of confluent cells. Although mechanisms that shape tissue sheets are increasingly understood, those which shape a volume of cells remain obscure. Here we show 3D mesenchymal cell intercalations, rather than cell divisions and biophysical tissue properties, are essential to shape the mandibular arch of the mouse embryo. Using a genetically encoded vinculin tension sensor, we show that cortical force oscillations promote these intercalations. Genetic loss and gain of function approaches show that *Wnt5a* functions as a spatial cue to coordinate cell polarity with cytoskeletal oscillation. YAP/TAZ and PIEZO1 serve as downstream effectors of *Wnt5a*-mediated actomyosin bias and cytosolic calcium transients, respectively, to ensure appropriate tissue form during growth. Our data support oriented 3D cell neighbour exchange as a conserved mechanism driving volumetric morphogenesis.

## INTRODUCTION

Morphogenesis refers to the process of shaping tissue during development, the reproducible nature of which is essential for appropriate pattern formation and function. Most of the recognised principles of morphogenesis concern mechanisms that shape sheets of embryonic tissue^1–13^. In particular, exchange of cell neighbours is central to tissue shape change in two dimensions (2D) and involves a limited number of transient multicellular configurations including tetrads (T1 exchange) and rosettes. Actomyosin contractions generate forces that promote cell neighbour exchanges and are oriented, in part, by physical properties of tissue that may be anisotropic in nature^14–16^. Some of these principles have been extended to curved epithelial sheets by combining empirical and theoretical approaches^17–19^. However, it remains unclear whether similar mechanisms apply to mesenchymal tissues, in part because mesenchymal tissues have been regarded as less confluent due to the presence of abundant extracellular matrix in some contexts and to potentially less adherent cell-cell junctions^20^. Changes in the viscoelastic properties of tissue are also associated with, and may partly drive, morphogenetic movements^21–24^, although the relationship between cellular and tissue scale properties remains unclear and may be context-dependent.

Multiple organ primordia such as the branchial arches, limb buds and genital tubercle are composed of an internal bulk layer of mesenchyme. In models of multilayered vertebrate tissues such as the frog gastrula, mechanisms of morphogenesis include amoeboid endodermal cell movements^25^ and mesodermal cell intercalations through junctional remodelling, though the latter takes place in a sheetlike manner^26,27^. Another example is elongation of the rod-like skeletal anlage in the vertebrate limb that is attributable to a highly structured columnar arrangement of chondrocytes embedded within abundant extracellular matrix and to oriented rearrangement of nascent daughter cells^28,29^. In contrast to these examples, very little is known about how more-or-less isotropic volumes of confluent cell organise to generate morphogenetic movements. For example, although the directional nature of mesodermal cell movements from the lateral plate to the limb bud has been demonstrated^30–34^, the cellular basis of those movements, and of volumetric morphogenesis in general, remain largely uncharacterised.

Early branchial arches are composed of a core volume of mesenchyme that is surrounded by a single cell layer epithelium. Although the neural crest^35–37^ and cranial mesodermal^38^ origins of branchial arch mesenchyme are well recognised, mechanisms by which the structure grows outward and acquires shape are less clear. Loss-of-function approaches that were intended to define the roles of various signalling pathways identified cellular processes that are relevant to branchial arch morphogenesis, such as cell survival^39–43^, cell proliferation^44,45^ and cell migration^46–48^. Those studies generated important insights, but were not intended to explain how the branchial arches acquire shape.

Craniofacial anomalies are common among birth defects. The mandibular portion of the first branchial arch generates multiple facial structures including the lower jaw, and some syndromes are associated with structurally common features of malformation such as a short (front to back) and broad (side to side) mandible. One of these, autosomal-dominant Robinow syndrome, is caused by mutations in components of the noncanonical WNT pathway including *WNT5A* and *ROR2*^49–52^. *Wnt5a* mutant mice that phenocopy many of the features of the human syndrome exhibit a short and bulbous mandibular arch^50,53^, though the cellular and physical basis of the malformation is not clear.

Here we study the mandibular arch as a model of two distinct modes of 3D morphogenesis. We show that cell division and physical tissue properties are important for growth but do not sufficiently explain how the arch primordium acquires a narrow mid-portion and a bulbous distal portion. Our data support a model in which 3D mesenchymal cell intercalations narrow and elongate the mid-portion. Relatively high amplitude cortical force oscillations promote cell intercalations in a *Wnt5a*– and *Piezo1*-dependent manner, implying these regulators spatially fine tune physical cell behaviours.

## RESULTS

### Spatial distributions of cell division frequency and viscoelastic tissue properties are insufficient to explain mandibular arch shape

The first branchial arch buds at the ventral aspect of the midbrain-hindbrain boundary in the mouse embryo. As shown by optical projection tomography (OPT), the mandibular portion of the first branchial arch remodels from a rod-like structure at the 19 somite stage (~E9.0) to form a narrow central ‘waist’ and a distal bulbous region between somite stages 21-28 (~E9.25-E9.75; Fig. 1A). In this study, we focused on the shape change between 19 and 21 somite stages (a ~4 h period) (Supplementary Mov. 1, 2) to understand how the waist and bulbous regions become defined.

**Figure 1.**
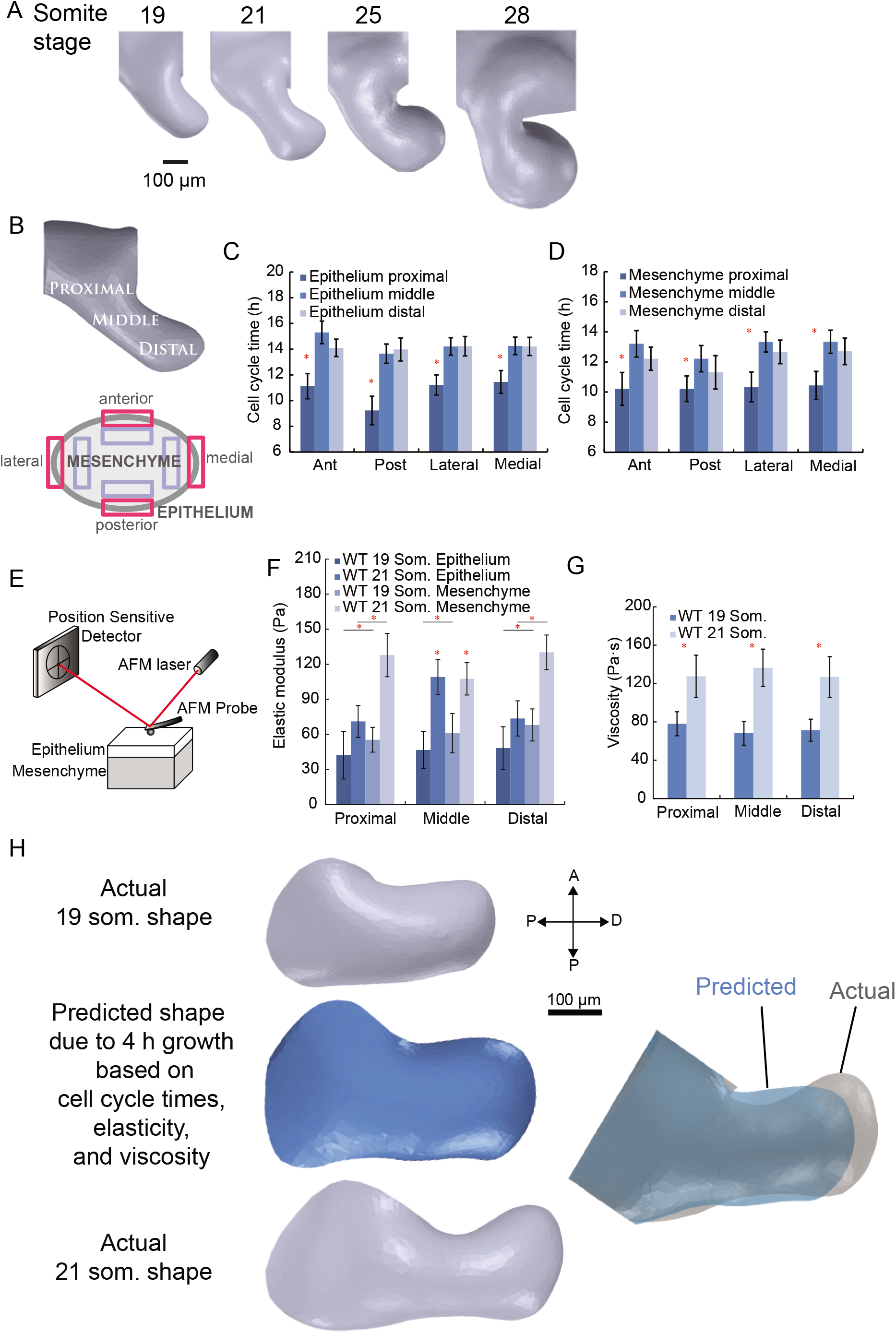
Spatial distributions of cell cycle time, elastic modulus and viscosity are insufficient to explain mandibular arch shape. **A** Sagittal OPT renderings of the right mandibular arch in the mouse embryo at different stages. **B** Cell cycle time was measured in 24 adjacent regions (4 epithelial and 4 mesenchymal regions each within proximal, middle and distal arch) of the 19 somite stage mandibular arch. **C**, **D** Spatial variation of epithelial (C) and mesenchymal (D) cell cycle times in the mandibular arch. Cell division was more rapid in the proximal region for both epithelial and mesenchymal layers; n=3 embryos at 20 somite stage, 15-35 cells examined for each of 12 epithelial regions per embryo, 50-75 cells examined for each of 12 mesenchymal regions per embryo; asterisks denote p<0.05, error bars denote standard error of the mean (s.e.m.). **E** Tissue indentation by AFM was employed to measure properties of intact, live mouse embryos. **F** Elastic (Young’s) modulus (stiffness) of epithelium and mesenchyme. **G** Viscosity of whole tissue in proximal, middle and distal regions of the mandibular arch at 19 and 21 somite stages. For F and G, 15 separate sites in each proximal, middle and distal region were indented in triplicate (45 measurements per region) per embryo; n=3 embryos, asterisks denote p≤0.05, error bars denote standard deviation. **H** Finite element simulation of 4 h of growth beginning from the actual 19 somite stage mandibular arch shape to predict 21 somite stage shape. The model incorporated experimentally measured spatial variation of cell cycle time, elasticity and viscosity. Simulated growth (in blue) results in an arch that is shorter and broader than the actual 21 somite stage arch.

To examine whether spatial variation in the frequency of cell division influences tissue shape, we measured cell cycle times using dual pulse with two thymidine analogues (5-Bromo-2’-deoxyuridine/iododeoxyuridine (BrdU/IdU))^54^. Each of the epithelial and mesenchymal tissue layers were arbitrarily divided into twelve spatial regions within which mean cell cycle times were calculated (Fig. 1B, Supplementary Fig. 1A). Cell division was most rapid in the proximal third of the mandibular arch adjacent to the face but there was little difference between the middle and distal thirds (Fig. 1C,D), implying this parameter does not intuitively explain differences in waist/bulbous morphology.

Physical tissue properties profoundly influence morphogenesis. Using atomic force microscopy (AFM) to measure elasticity (Fig. 1E, Supplementary Fig. 1B), we performed shallow (1 μm) indentation of the ~12 μm thick epithelium to exclude substrate effects of the underlying mesenchyme (Supplementary Fig. 1C) and found that waist epithelium becomes stiffer relative to more proximal and distal flanking regions between somite stages 19 and 21 (Fig. 1F). Mesenchymal stiffness was measured using deep (7-9 μm) indentation (Supplementary Fig. 1D) and decoupled from the influence of the epithelium (see Methods). In contrast to the epithelium, mesenchyme was least stiff in the middle region (Fig. 1F). Under shallow indentation, the arch tissue behaved as an elastic material (Supplementary Fig. 1C), whereas deep probe indentation and retraction revealed evidence of hysteresis, or dissipation of energy, an indication of viscoelasticity (Supplementary Fig. 1D). Combined (epithelial and mesenchymal) tissue viscosity that was quantified using different rates (5, 10, and 15 μm/s) of indentation (Supplementary Fig. 1E) also increased over developmental time but did not vary spatially throughout the arch (Fig. 1G). An unbiased model would be useful to evaluate the potential morphogenetic relevance of these data.

To integrate the influence of cell divisions and physical properties on tissue shape, we constructed a 3D, two tissue layer finite element model of the 19 somite stage mandibular arch that was based on OPT-derived tissue topography (Supplementary Fig. 1F, G), an approach conceptually similar to that previously reported for limb bud shape^30^ but with the addition of empirical physical properties. Based on prior work, we initially hypothesised that disproportionately increased stiffness of the middle epithelium might resist hoop stress due to growth of the mesenchyme to yield a growth process reminiscent of directed dilation^55,56^. However, simulated deformation predicted an inappropriately short and wide arch (Fig. 1H, Supplementary Mov. 3). We reasoned that the inadequacy of this continuum model might reflect the fact that it did not account for defined cell rearrangements.

### Distinct cellular parameters characterise two regions of the mandibular arch

In confluent 2D epithelia, defined neighbour exchange processes are essential for tissue remodelling. We noticed that the mesenchymal cells within the mandibular arch exhibited key characteristics of mouse epithelial cells that rearrange through neighbour exchange, such as confluence, abundant expression of cell-cell junction proteins (N-cadherin and desmoglein), and protrusive activity (Supplementary Fig. 2A-C)^14,57^.

Since biological parameters that are relevant to the possibility of 3D cell intercalation have not been well defined, we explored ideas from the physical sciences that are hypothetically relevant to morphogenesis. We performed live lightsheet microscopy of intact mouse embryos by combining transgenic CAG::H2B-GFP and mTmG reporters to highlight nuclei in green and cell membranes in red (Supplementary Mov. 4, 5). As in an unstable foam^58,59^, mesenchymal cells exhibited a broad distribution of cell faces (*7-14*) with relatively few neighbours in the mid-portion of the mandibular arch (Fig. 2A,B, Supplementary Fig. 2D, Supplementary Mov. 6), suggesting cells in that region are furthest from equilibrium and most likely to exchange neighbours. To estimate cell shapes, we segmented nuclei in 3D to act as centroids for Voronoi tessellation (Fig. 2C). According to a recent model, there is a relationship between the rigidity of confluent 3D tissues and the ratio of cellular surface area to volume^60^. The cell shape parameter surface area/volume^2/3^ (S/V^2/3^) varied along the proximodistal axis of the mandibular arch with the highest values observed in the middle region (Fig. 2D,E), that is consistent with a relatively more ‘liquid’ behaviour compared to proximal and distal regions. These parameters predict that middle region cells should preferentially engage in neighbour exchange.

**Figure 2.**
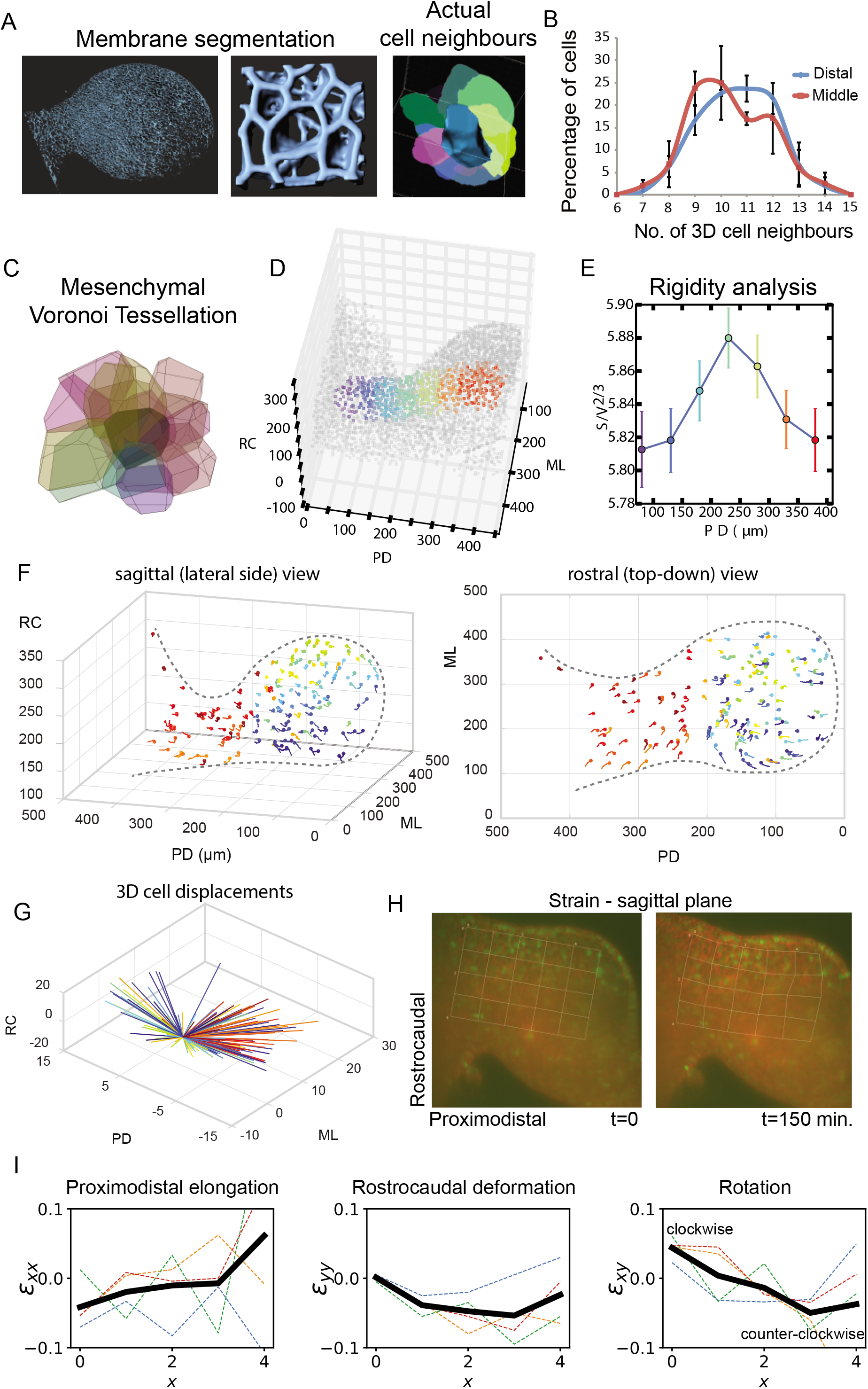
Two distinct patterns of growth characterise the mandibular arch. **A** 3D renderings of cell membranes labelled by *mTmG* based on live light sheet microscopy. Whole arch (left) and local cell neighbour relations (middle and right) are shown. **B** Distribution of numbers of cell neighbours in middle (red curve) and distal (blue curves) mandibular arch (n=2 embryos, 202 middle cells and 244 distal cells examined, p<0.05, chi-squared). **C** Voronoi tesselation of mesenchymal nuclei to estimate cell shapes. **D**, **E** Colour-coded distribution of cell shape index (S/V^2/3^) that correlates positively with liquid-like tissue phase (greatest in the middle region). **F** 4D tracks of a subset of mandibular arch cells are shown in two orthogonal views. Relatively directional tracks that were oriented distalward characterised the waist region, whereas short and tightly curved tracks characterise the bulbous region. **G** 3D dandelion plot of colour-coded trajectories of cells at the start and end of a movie. Waist cells (red/orange) move predominantly outward whereas bulbous cells move in a relatively radial fashion. **H** Strain illustrated as deformation of a sagittal plane grid during a 150 min. movie. Nodes remain fixed to the same positions throughout the movie. I The proximodistal axis extended (reflected by the positive ε_xx_slope), while the midportion of the arch converged along the rostrocaudal axis (reflected by negative values of the ε_yy_ curve). Convergence of midportion tissue combined with expansion of distal tissue results in clockwise (positive ε_xy_ values) and counter-clockwise (negative ε_xy_ values) rotational deformation of adjacent regions, respectively (as reflected by the downward ε_xy_ slope). Coloured lines correspond to components of the strain tensor for each square along the rostroacaudal axis. The black line is the average of rostrocaudal strain tensors as it varied along the proximodistal axis.

To identify the large-scale pattern of tissue displacement, a subset of fluorescently labelled nuclei were tracked in 4D after accounting for embryo drift by marking multiple fluorescent beads that we co-embedded with the embryo in an agarose cylinder. Tissue growth was relatively longitudinal in the middle region compared to the distal region (Fig. 2F,G, Supplementary Mov. 7,8). The cumulative distribution function (CDF) of persistence time (the extent to which cells move in accordance with their recent past^61,62^) exhibited a greater slope for cells in the middle compared to those in the distal region, indicating the former are more directionally consistent over time (Supplementary Fig. 2E–H, Supplementary Mov. 9, 10). Therefore, distinct morphogenetic movements characterise the middle and distal regions of the arch.

### Cell intercalations in 3D promote volumetric convergent extension

Cell displacements, by themselves, are not evidence of active cell movements since the tissue they inhabit expands due to cell divisions. Analysis of tissue strain indicated that proximodistal elongation of the arch was accompanied by compression along the short rostrocaudal axis of the midportion (Fig. 2H,I, Supplementary Mov. 11), an observation that can potentially be explained by convergent cell movements. To test for this possibility, we first examined epithelial cells at higher resolution using live confocal time-lapse imaging. In the middle epithelium, although the alignment of daughter cells was biased along the short rostrocaudal axis, T1 exchange events tended to shorten the rostrocaudal axis and elongate the proximodistal axis of growth. Cell rearrangements were less oriented in the distal region (Fig. 3A,B, Supplementary Fig. 3A), consistent with bulbous growth.

**Figure 3.**
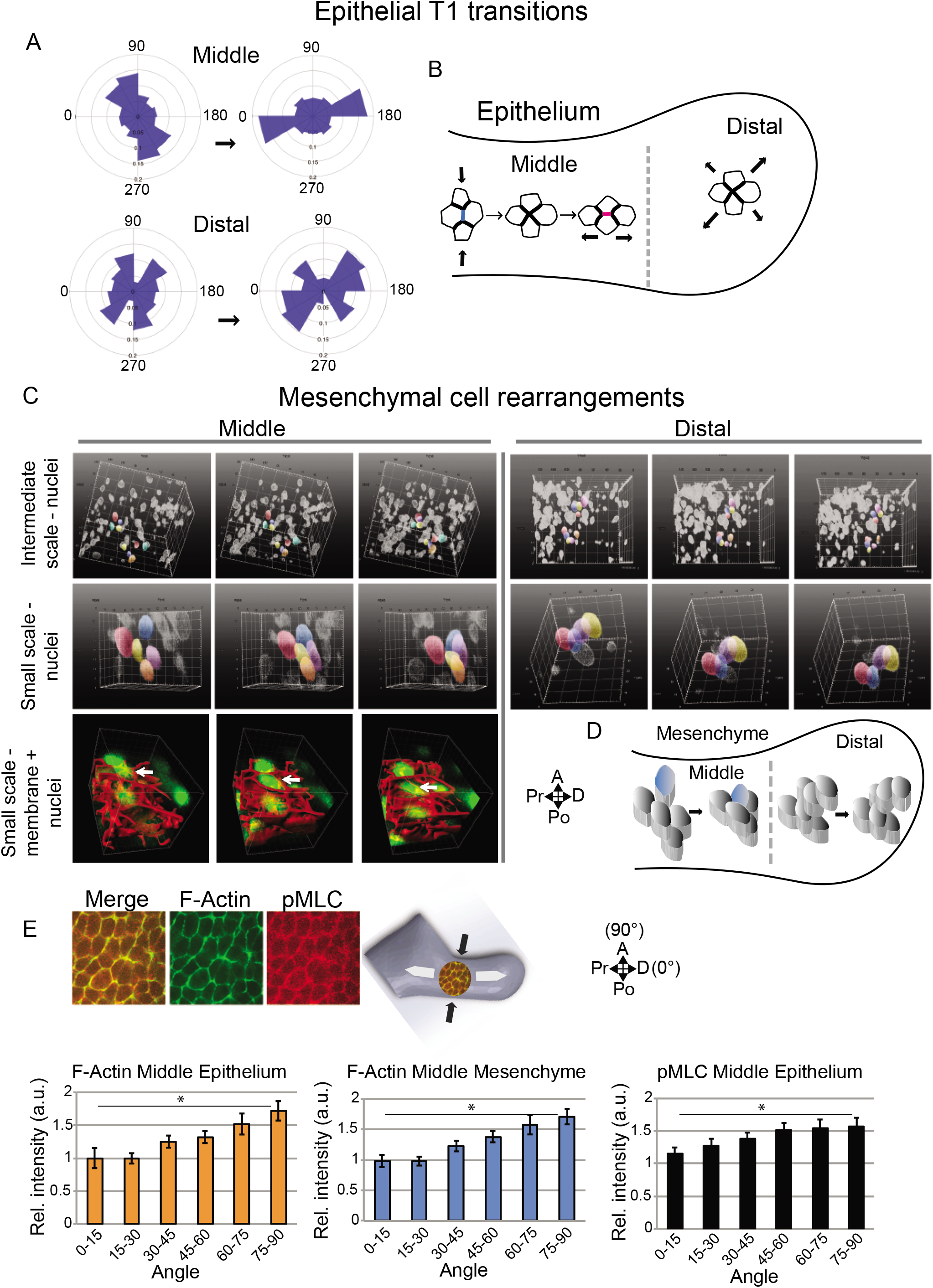
Epithelial and mesenchymal cell rearrangements converge and extend the midportion of the mandibular arch. **A** Orientation of epithelial tetrads during T1 transitions at formation to resolution stages (separated by an arrow). **B** Schematic representation of predominant orientation of epithelial T1 transitions in middle and distal regions. **C** Time series of volumes of mesenchymal cells of dual *H2B-GFP;mTmG* transgenic embryos visualised by light sheet microscopy at intermediate and high magnification. Select nuclei are coloured to show tissue and cell convergence at intermediate and small scales occurs in the middle, but not distal, region. **D** Schematic representation of oriented mesenchymal cell intercalations transverse to the axis of elongation in the middle region. **E** In the mid-portion of the arch, F-actin and phosphomyosin light chain (pMLC) were biased along proximal and distal epithelial and mesenchymal cell interfaces which is parallel to the rostrocaudal axis and to the direction of cell intercalations. The angular distribution of immunostain fluorescence intensity for epithelial (n=4 embryos) and mesenchymal (n=4 embryos) F-actin and epithelial phosphomyosin light chain (pMLC) (n=5 embryos) relative to the arch long axis that was designated as 0° was quantified in the middle region using SIESTA. Asterisks denote p<0.05, error bars denote s.e.m.

For mesenchyme, live light sheet movies at intermediate scale (~100 cells) revealed that cells in the middle region converged centripetally relative to one another as tissue growed longitudinally (Fig. 3C, Supplementary Mov. 12). At small scales (~10 cells), the intercalation of a single cell into a nest of 5-6 others was the smallest multicellular unit in which 3D neighbour exchange was observed (Fig. 3C, Supplementary Mov. 13-15). This configuration is analogous to 3D T1 exchange in an unstable foam^59^. Tracking of nuclear centroids undergoing intercalation confirmed that small groups of cells converged in the axial plane as the tissue extended distalward (Supplementary Fig. 3B). In contrast, more subtle intercellular adjustments characterised the distal region (Fig. 3C, Supplementary Fig. 3C, Supplementary Mov. 16). These observations support the concept that regional differences in cell intercalation correlate with large-scale growth patterns and that volumetric convergent extension elongates the middle region (Fig. 3D).

Cortical actomyosin can orient forces that drive cell intercalation in epithelia and in mesoderm^26,63,64^. F-actin and phospho (p)-myosin light chain (pMLC) were biased parallel to the short, transverse axis among epithelial and mesenchymal cells in the middle waist, but not proximal and distal regions, of the arch (Fig. 3E, Supplementary Fig. 3D). This bias likely reflects the tissue stress pattern^14^ and is consistent with the axis of intercellular movements observed in the waist region, but itself does not explain why cells intercalate.

### Oscillatory amplitude of cortical tension correlates with mesenchymal cell intercalation

Cortical tension is a key parameter that regulates cell sorting and intercalation during development^65–67^. Local actomyosin abundance or cell interface length observations have been used as a proxy for cortical tension, but may not reflect either actual forces or their dynamic fluctuations. To directly measure cortical tension, we knocked-in a conditional FRET (Förster resonance energy transfer)-based vinculin tension sensor (VinTS), for which fluorescence lifetime in nanoseconds (ns) is proportional to force in piconewtons (pN)^68^, into the mouse *Rosa* locus. We also generated two control knock-in strains that should exhibit maximal (donor only VinTFP – no FRET), and minimal (vinculin tailless VinTL – maximal FRET) fluorescence lifetime, respectively (Fig. 4A).

**Figure 4.**
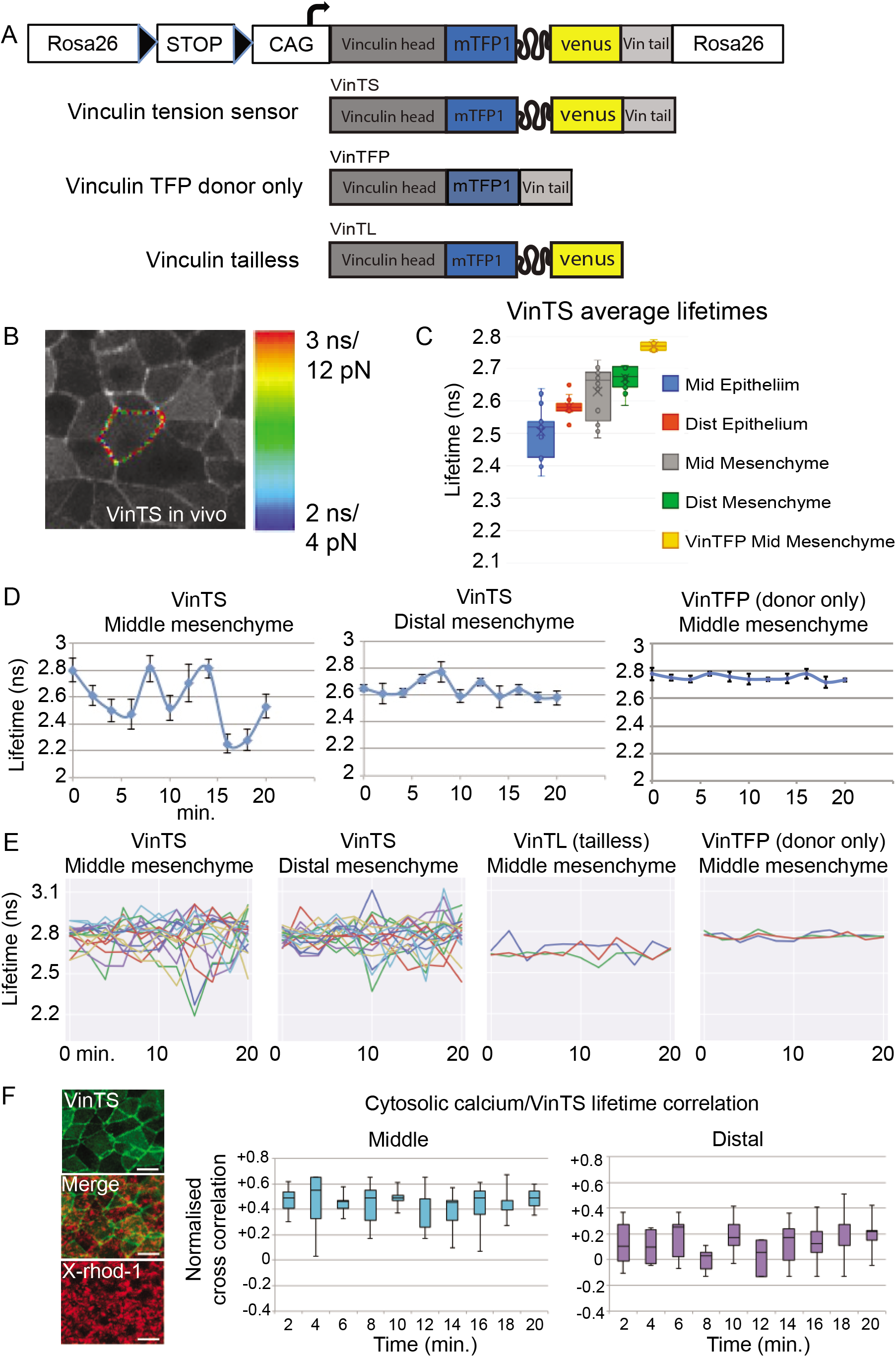
Vinculin force oscillations distinguish middle and distal regions of the mandibular arch. **A** Conditional *Rosa26* knock-in mouse strains: full length vinculin tension sensor (VinTS), TFP (FRET donor) only control (VinTFP), vinculin tailless control (VinTL). **B** Tension sensor expression among epithelial cells in the mandibular arch with one cell cortex highlighted as region of interest. Colour scale shows range of lifetime (in nanonseconds, ns) and corresponding force values (in picoNewtons, pN). **C** Individual cell fluorescence lifetime values in middle (mid) and distal (dist) epithelium and mesenchyme of the mandibular arch. Boxplots show mean (x), median (—), central quartiles (coloured box), and range (transverse end bars); n=15 cells per region in each of 3 embryos. **D** Representative vinculin force curves of individual cells in middle and distal regions, and donor only control. Lifetime readings were taken at two minute intervals, error bars denote s.e.m. **E** Analysis of multiple vinculin force curves revealed that the sample variance of lifetime values, a measure of amplitude, was greater among middle (0.0201 ns) versus distal (0.0132 ns) mesenchymal cells (p=0.03, t-test). Mean lifetime variance was lower among middle mesenchymal cells of control VinTL (0.0028 ns, p=0.02) and VinTFP (0.0006 ns, p=0.01) strains. **F** Correlations of VinTS fluorescence lifetime and calcium reporter X-rhod-1 fluctuation (defined the percentage of X-rhod-1 AM staining area for each cell at each time point) in middle and distal arch regions over time.

All three knock-in constructs were expressed robustly in the appropriate cortical domain *in vitro* (Supplementary Fig. 4A) and *in vivo* (Fig. 4B, Supplementary Mov. 17). Measurements of the dynamic range, floor and ceiling lifetime values of the three strains were undertaken in ES cell colonies and embryoid bodies prior to their evaluation in the mouse embryo under conditions that we previously optimised for live imaging^14,32^.

Single cell cortices were outlined as regions of interest to measure lifetime. As expected, the donor-only VinTFP construct reported the longest lifetime with a narrow standard deviation and VinTL exhibited short lifetime values. In embryoid body cells and among differentiated beating cardiomyocytes, the range and standard deviation of the full length VinTS lifetime values was greater than that of either control suggesting the reporter was responding dynamically to cell contractions (Supplementary Fig. 4B, C). Among epithelial and mesenchymal cells of the mandibular arch, VinTS reported greater lifetime range and standard deviation than either control strain indicating robust dynamic capacity *in vivo* (Supplementary Fig. 4D). Treatment of live transgenic embryos with the Rock inhibitor Y27632 or the actin polymerisation inhibitor Cytochalasin D lowered average VinTS lifetime values to floor levels for this construct and dampened standard deviation values measured in the mandibular arch (Supplementary Fig. 4E, F). These data indicate the VinTS sensor reports forces attributable to actomyosin contraction *in vivo*.

For both epithelial and mesenchymal tissue layers, cells in the middle region exhibited lower cytoskeletal tension along with greater variability compared to the corresponding distal region (Fig. 4C). Since high cortical tension generally reflects relatively spherical cell shapes, these differences are consistent with face numbers we documented and support the concept that cells in the middle region are more likely to intercalate. Time-lapse analysis with two-minute intervals revealed an oscillatory pattern of fluorescence lifetime fluctuation among individual cell cortices using the VinTS reporter. In support of a previous theoretical assertion that membrane fluctuation is necessary to overcome the energy barrier for cell intercalation^23,24^, middle region mesenchymal cells exhibited a significantly greater fluctuation amplitude compared to those in the distal region (Fig. 4D,E).

Transient increases in cytosolic Ca^2+^ ion concentration are required for contractile cortical pulses^69,70^. We applied a Ca^2+^ indicator (X-Rhod-1 or Fluo8) to intact mouse embryos and quantified the proportion of X-Rhod-1 fluorescence to total cell area (marked by VinTS). Cytosolic Ca^2+^ fluctuated in an asynchronous fashion among waist mesenchymal cells and was relatively dampened among distal cells (Supplementary Mov.18,19, Supplementary Fig. 4G) (fig. S6G). Ca^2+^ concentration correlated with VinTS fluorescence lifetime in the middle, but not distal, region of the arch (Fig. 4F), suggesting Ca^2+^ fluctuation promotes cortical oscillation.

### *Wnt5a* acts upstream of YAP/TAZ and PIEZO1 to regulate cortical polarity and oscillation

We observed that *Wnt5a^-/-^* mouse mutants, which substantially phenocopy Robinow syndrome^50,53^, exhibit a proximodistally short and mediolaterally broad mandibular arch (Fig. 5A, Supplementary Mov. 20, 21). Cell cycle times in *Wnt5a^-/-^* embryos were similar to those of WT embryos (Supplementary Fig. 5A, B, compare with Fig. 1C, D), although stiffness and viscosity did not increase to the same degree as in WT tissue, especially in the middle region (Supplementary Fig. 5C, D, compare with Fig. 1F, G). Finite element simulation of mandibular arch growth more closely matched actual mutant arch shape as compared to the WT simulation (Fig. 5B,C compare with Fig. 1H, Supplementary Mov. 22), suggesting that the influence of cell rearrangements might be less relevant to growth in the *Wnt5a^-/-^* mutant. Consistent with that concept, mutant cells exhibited directionally less persistent movements that were reflected by a lower slope for cumulative distribution function (CDF) for both persistence time and angle from the mean compared to cells in WT embryos (Fig. 5D, E).

**Figure 5.**
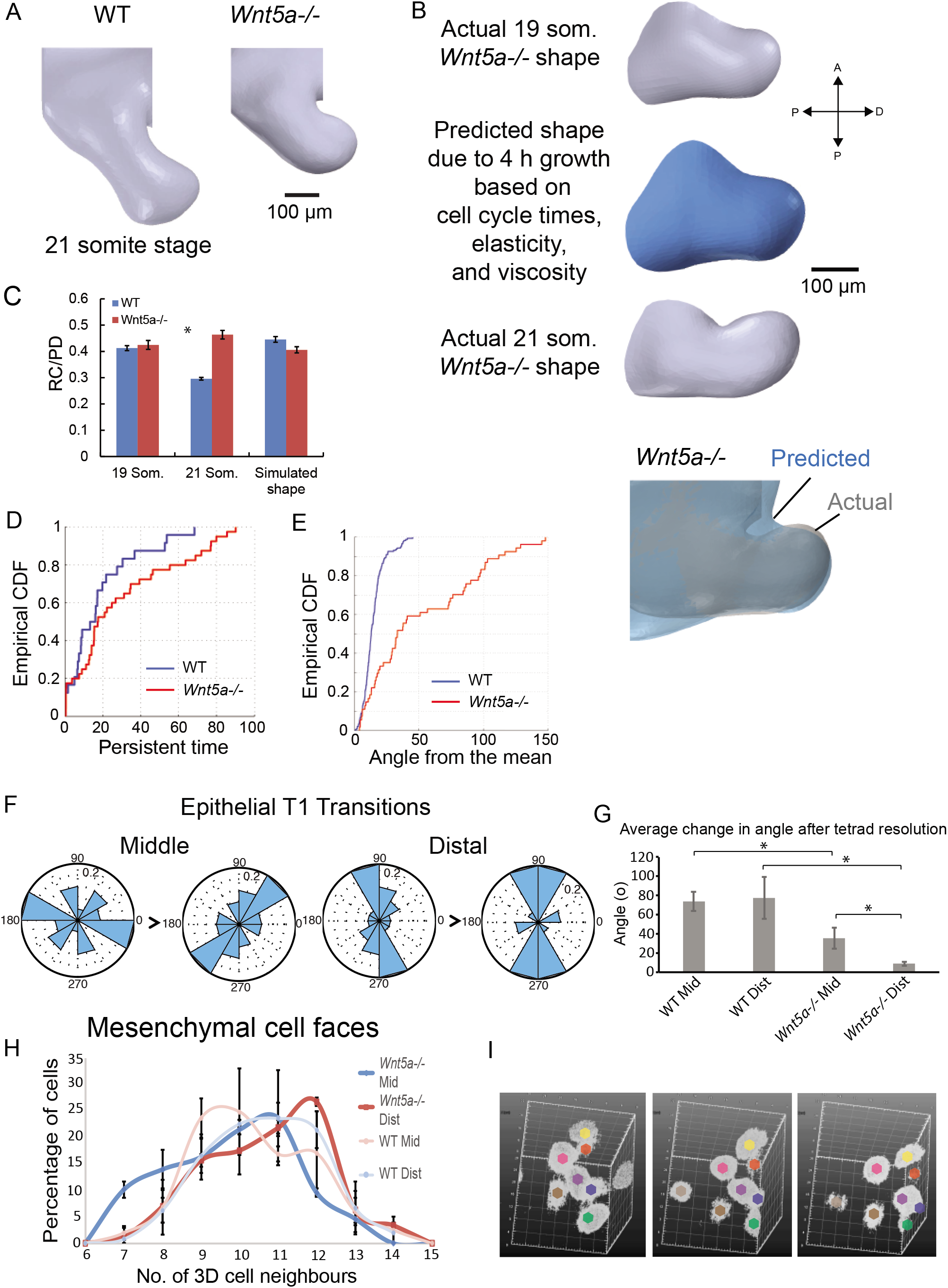
Deficient cell rearrangements in *Wnt5a^-/-^* mutants. **A** The *Wnt5a^-/-^* mutant mandibular arch is comparatively short and broad by OPT. **B** A finite element model incorporating the spatial variation of *Wnt5a^-/-^* mutant cell cycle time, Young’s modulus and viscosity was employed to simulate 4 h of growth starting from the actual 19 somite stage *Wnt5a’ ^/^’* mutant mandibular arch shape compared to the actual 21 somite stage *Wnt5a^-/-^* mutant arch. **C** Ratio of rostrocaudal to proximodistal axis length (RC/PD) was comparatively better for finite element simulation of *Wnt5a^-/-^* mutant growth than for WT. **D**, **E** Nuclei of *H2B~GFP* transgenic embryos were tracked by 4D light sheet microscopy and, according to a random walk model, persistence of cell movements (persistent time, (B)) and direction (angle from the mean, (C)) were diminished in *Wnt5a* mutants. CDF: cumulative distribution function. **F** Epithelial T1 transitions in the *Wnt5a^-/-^* mutant mandibular arch did not tend to converge and extend the middle region as in WT (compare with Fig. 3B). **G** Angular change in long axis among resolving tetrads was diminished in *Wnt5a^-/-^* mutant epithelium (n=entire lateral epithelium in each of 3-5 embryos per condition, asterisks indicate p<0.05). **H** Distribution of mesenchymal cell face numbers was shifted rightward to higher values for *Wnt5a^-/-^* mutant mesenchyme (n=2 *Wnt5a^-/-^* embryos, 92 middle cells, 146 distal cells, error bars denote s.e.m.). I Representative group of mesenchymal cells in the mutant middle region that did not intercalate as in the WT middle region, but rather resembled subtle intercellular adjustments as in WT distal mesenchyme (compare with Fig. 3C).

In *Wnt5a^-/-^* mutants, 2D epithelial T1 exchanges were disoriented (Fig. 5F, G, compare with Fig. 3A), mesenchymal cells exhibited a rightward-shifted (more stable) distribution of cell neighbour numbers (Fig. 5H), and the frequency of mesenchymal intercalations was diminished in the middle region (Fig. 5I, Supplementary Fig. 5E, Supplementary Mov. 23, 24). Cortical F-actin, phosphomyosin light chain (pMLC), and VANGL2 were not biased along rostrocaudal cell interfaces in *Wnt5a^-/-^* mutants (Fig. 6A, Supplementary Fig. 6A-C). Moreover, the amplitude of cortical oscillations (Fig. 6B-D) and Ca^2+^ fluctuations (Fig. 6E, Supplementary Fig. 6D, Supplementary Mov. 25, 26) were diminished in the mutant middle arch. Together, these data imply that *Wnt5a* acts through the cell polarity and Ca^2+^ pathways to promote and orient cell intercalations.

**Figure 6.**
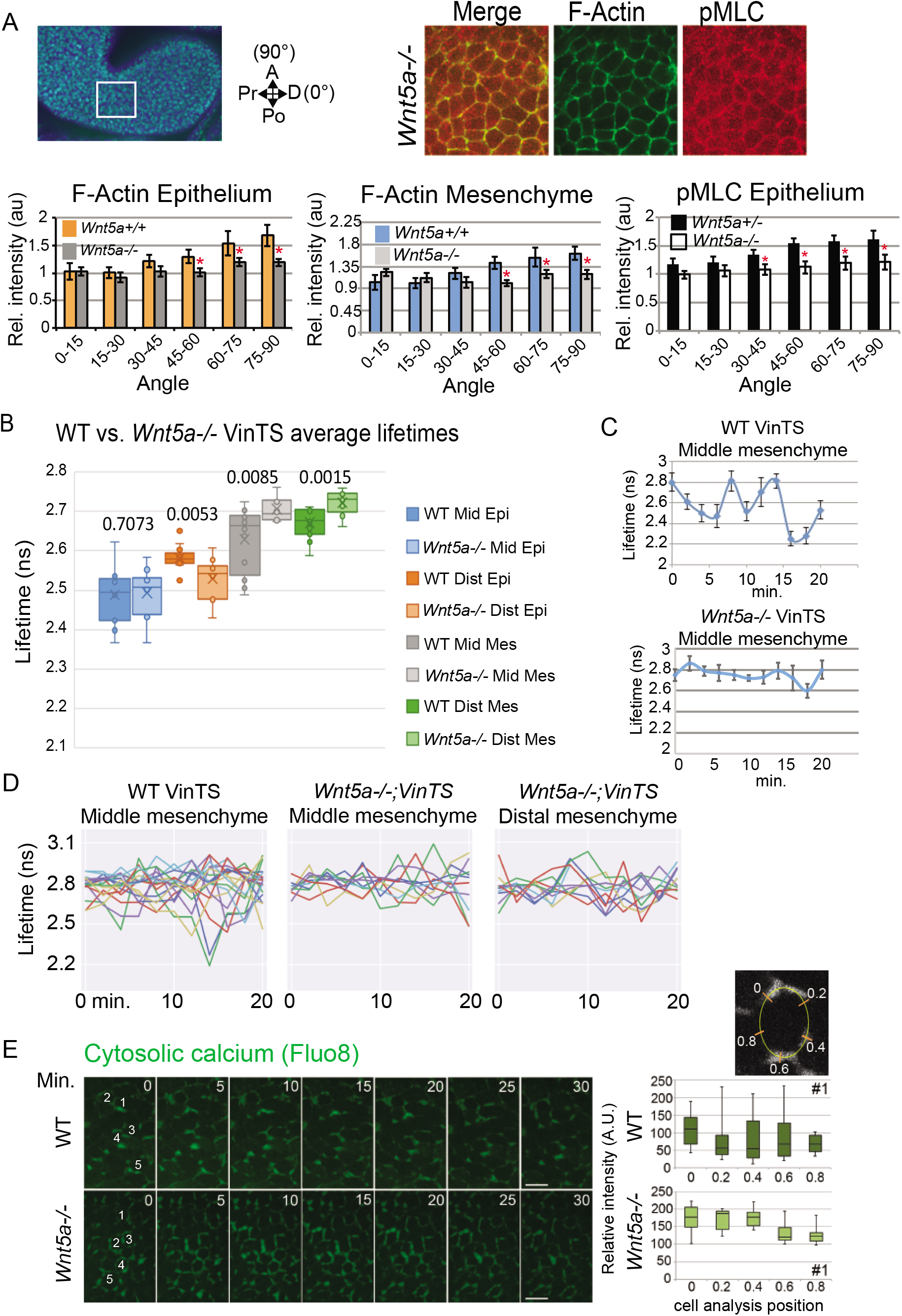
Actomyosin bias and vinculin tension were diminished in the *Wnt5a^-/-^* mutant mandibular arch. **A** Spatial distributions of F-actin and pMLC were not biased in 21 somite stage *Wnt5a^-/-^* mutants. The angular distribution of immunostain fluorescence intensity for epithelial (n=4 embryos) and mesenchymal (n=4 embryos), and epithelial pMLC (n=5 embryos) was quantified relative to the arch long axis that was designated 0° using SIESTA. Asterisks denote p<0.05, error bars denote s.e.m. **B** Average fluorescence lifetime values in middle (mid) and distal (dist) epithelium and mesenchyme of mandibular arch. Boxplots show mean (x), median (—), central quartiles (coloured box), and range (end bars); n=15 cells per region x 3 embryos. P values for pairwise WT/*Wnt5a^-/-^* mutant comparisons are given in the graph. Of note, middle mesenchymal lifetime range was diminished in *Wnt5a^-/-^* mutants and more closely resembled WT distal mesenchymal lifetime values. **C** Representative vinculin force curves of an individual *Wnt5a^-/-^* mutant cell in the middle region that lacks the amplitude observed for WT cells in same region. **D** Multiple vinculin force curves revealed that the variance (amplitude) of lifetime values for *Wnt5a^-/-^* mutant middle mesenchyme (0.0122 ns) was diminished relative to WT middle mesenchyme (0.0201 ns, p=0.04), but similar to WT distal mesenchyme (0.0132 ns, p=0.38). Lifetime variance was similar between WT (0.0132 ns) and *Wnt5a^-/-^* mutant (0.0096) distal mesenchyme (p=0.13). **E** Variation of cytosolic calcium concentration using Fluo8 applied to embryos was quantified at 5 positions per cell over time. Corresponds to Supplementary Fig. 6D.

The *Wnt5a* expression domain is biased distally and diminishes steeply within the middle region of the arch, but it was not clear whether it acts instructively or permissively to influence cortical behaviour. To test between these possibilities, we overexpressed a transgenic *Cre*-activated *Wnt5a* allele that was targeted to the endogenous *Ubiquitin-b* (*Ubb*) locus using *Sox2:Cre* to drive ubiquitous expression and diminish *Wnt5a* grandients (Fig. 7A). In *Sox2:Cre;Z/Wnt5a* embryos, the mandibular arch was short and broad (Fig. 7B, Supplementary Mov. 27), and lacked actomyosin polarity (Supplementary Fig. 7A), implying that the native *Wnt5a* expression domain provides a spatial cue for cortical organisation.

**Figure 7.**
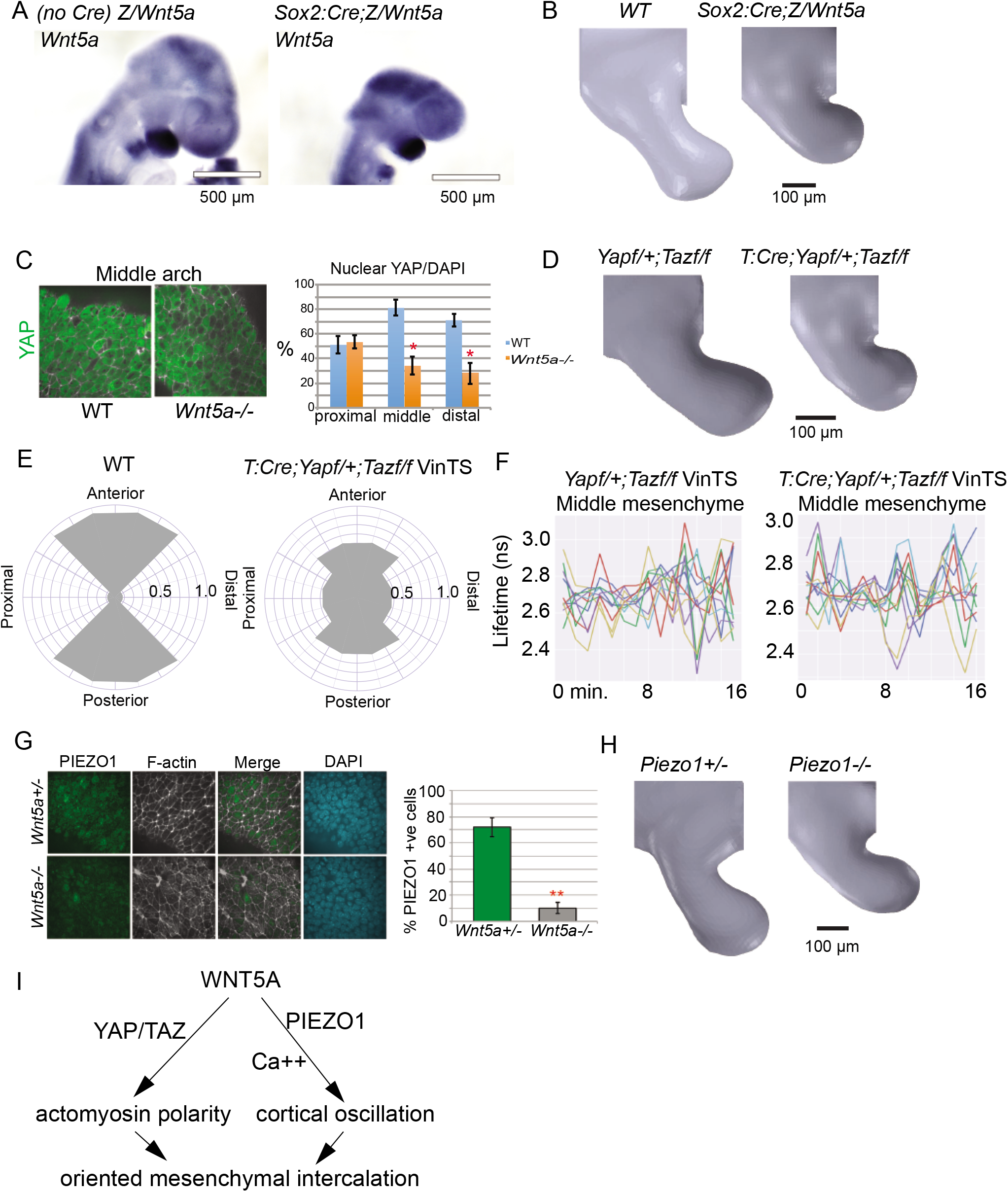
*Wnt5a* overexpression, *Yap/Taz* and *Piezo1* mutant analyses. **A** *Sox:Cre;Z/Wnt5a* embryos expressed *Wnt5a* beyond the normal expression domains by in situ hybridisation. **B** OPT showing a short and wide mandibular arch in a 21 somite *Sox:Cre;Z/Wnt5a* embryo. **C** Proportion of cells exhibiting nuclear YAP in the *Wnt5a^-/-^* mandibular arch. **D** OPT showing phenotype of *T:Cre;Yap^f/+^;Taz^f/f^* mandibular arch. **E** Orientation of mesenchymal cell intercalations in the sagittal plane of mandibular arch mid-regions of WT and *T:Cre;Yap^f/+^;Taz^f/f^* embryos. Intercalation angles were binned into one of 4 arcs of 90° each: anterior, posterior, proximal and distal (n for WT=48, n for mutants=42, where n=No. of groups of 5-7 cells). **F** Multiple vinculin force curves showed no difference in the variance (amplitude) of fluorescence lifetime values of *Yap^f/+^;Taz^f/f^* and *Tcre;Yap^f/+^;Taz^f/^* middle mesenchymal cells (p=0.185, n=16 cells in each of two 20-21 somite embryos per condition). **G** PIEZO1 immunostain intensity was diminished in the *Wnt5a^-/-^* mandibular arch. **H** The 21 somite *Piezo1^-/-^* mutant mandibular arch partially phenocopied that of *Wnt5a^-/-^* with a short and broad midportion. **I** YAP/TAZ and PIEZO1 partially mediate actomyosin bias and cortical oscillation amplitude, respectively, downstream of *Wnt5a* to orient and promote mesenchymal cell intercalations.

We examined the Hippo pathway since it has been implicated in craniofacial malformation^57,58^, is mechanoresponsive^71,72^, and is a downstream effector of noncanonical *WNT5A* signalling that promotes YAP/TAZ nuclear accumulation^73^. In *Wnt5a* mutants, nuclear localisation of YAP was indeed diminished among mesenchymal cells of the middle and distal region of the branchial arch (Fig. 7C). We used *T:Cre* or *Wnt1:Cre* to excise *Yap/Taz* in early mesodermal somitomeres (~E7.5) or in neural crest, respectively, both of which contribute to branchial arch mesenchyme. Although deletion of all four *Yap/Taz* alleles caused lethality prior to branchial arch development, removal of any three *Yap* and *Taz* alleles with either Cre largely phenocopied *Wnt5a* null mutants with respect to mandibular shape (Fig. 7D). Loss of the upstream Hippo regulator *Fat4* had no impact on mandibular arch size or shape (not shown), suggesting the possibility that YAP/TAZ act primarily downstream of *WNT5A* in this context. Unlike in *Wnt5a* mutants, intercellular movements were abundant but disoriented from their orderly convergent intercalations among *Yap/Taz* mutants (Fig. 7E, Supplementary Mov. 28). Consistent with this finding, the mandibular arch in *Yap/Taz* mutants exhibited near normal cortical force amplitudes (Fig. 7F) in association with disrupted actomyosin polarity (Supplementary Fig. 7B). These data imply that cortical polarity is regulated independent of cortical oscillation, and is regulated by *Yap/Taz* in concert with or downstream of *Wnt5a*.

To further test whether cytosolic Ca^2+^ transients contribute to mandibular arch morphogenesis using another genetic perturbation, we examined *Piezo1* which encodes a mechanosensitive ion channel^74^. As expected, Ca^2+^ fluctuation was diminished in *Piezo1^-/-^* mutants (Supplementary Fig. 7C, Supplementary Mov. 29). PIEZO1 immunostain intensity was diminished in *Wnt5a^-/-^* mutants (Fig. 7G – antibody specificity was confirmed by its absence in *Piezo1* mutants – not shown), indicating its expression or stability is regulated by *Wnt5a*. The mandibular arch of *Piezo1^-/-^* embryos was misshapen similar to that of *Wnt5a^-/-^* mutants (Fig 7H, Supplementary Mov. 30), suggesting cell intercalations are disrupted. Together, these findings implicate YAP/TAZ and PIEZO1 as downstream effectors of *WNT5A*-mediated cortical polarity and oscillation, respectively (Fig. 7I).

## DISCUSSION

Our findings suggest that cell intercalations shape a volume of confluent cells, and that basic modes of 2D cell rearrangement, such as T1 exchange, have 3D counterparts previously observed in foams that remodel mesenchyme. Step-wise evolution of cell intercalation capacity from in-plane among diploblasts^75^, to out-of-plane among triploblasts^76^, to within a volume of cell neighbours may have facilitated the radiation of increasingly complex body plans among Bilateria and vertebrates.

By acting partly in the same pathway, *Wnt5a, Yap/Taz* and *Piezo1* transform biochemical signals to mechanical outputs that permit cell intercalation. In particular, oscillatory contractions of the cytoskeleton that have been observed in association with multiple types of invertebrate and vertebrate cell movements^9,63,77–83^ are likely essential for overcoming an energy barrier for cell intercalation^23,24^. The spatial coordination of cell polarity with cortical oscillation by *Wnt5a* upstream of YAP/TAZ and PIEZO1 effectors ensures that cell intercalations are oriented appropriately.

Explanations for the oscillatory nature of contractions that have been put forward include cell-extrinsic^84–86^ and cell-intrinsic mechanisms^87,88 69,89,90^, including Ca^2+^ flux^70,79^. In *Drosophila*, ion channel function is required to drive Ca^2+^ fluctuation and periodic cell contraction^70^, whereas in the mouse embryo, *Piezo1* partially fulfills this function. In our system, feedback between positive and negative regulators of Ca^2+^ influx, such as the noncanonical Wnt pathway and Ca^2+^-dependent proteases like calpain-2, respectively, have the potential to regulate oscillation amplitude.

Gradual stiffening of mandibular arch tissue that we observed is conceptually similar to what has been observed over the course of dorsal closure due to cell contractions in the amnioserosa of *Drosophila*^91^. The solidification of tissue after early morphogenetic movements establish primordial structures implies that a different suite of cell and tissue properties, possibly related to extracellular matrix and differentiating tissues, may regulate morphogenesis of solid organs at later stages. Combining increasingly accurate biophysical approaches with genetics will likely help to define morphogenesis pathways, including those that coordinate cell polarity with physical properties.

## ACKNOWLEDGEMENTS

We thank Phil Smallwood and Jeremy Nathans for the *Z/Wnt5a* mouse strain, Matthias Merkel, Lisa Manning and Rodrigo Fernandez-Gonzalez for helpful discussions and reviews of the manuscript. This study was supported by CIHR (MOP 126115) to SH and by Canada Research Chairs to YS.

## AUTHOR CONTRIBUTIONS

Experimental design: HT, MZ, KL, RA, YS, SH

Cell cycle times, immunostains, actomyosin polarity, Ca^++^ fluctuation, mutant analyses: HT

Atomic force microscopy, finite element modelling: MZ, XW, YS 4D cell tracking: OW

Vinculin tension sensor knock-in and measurements in vivo, light sheet imaging: KL

Vinculin tension sensor ceiling/floor assessments in vitro/in vivo: KL, NH, WL, MV, DH

Vinculin tension sensor anisotropy, group effects: KF, CP

Random walk model: XX, HH

Strain and rigidity analyses: MS, HH

Epithelial cell rearrangement analysis: XXC, MW

Mesenchymal cell rearrangement analysis: SH

Generation of Wnt5a overexpressing transgenics: EK, H-YH

Optical projection tomography: SS, MH

VANGL2 evaluation: HT, JF, RA

Vinculin tension sensor evaluation and advice: AD

Mutant analyses: HT, KL, MZ, JW

Biological question, manuscript preparation: SH

## METHODS

### Mouse strains

Analysis was performed using the following mouse strains: *CAG::H2G’GFP*^92^ (Jackson Laboratory: B6.Cg-Tg(HIST1H2BB/EGFP1Pa/J), *mTmG*^93^ (Jackson Laboratory: Gt(ROSA)26Sortm4(ACTB-tdTomato-EGFP)Luo/J)), *CAG::myr-Venus^94^*, full length vinculin tension sensor (*VinTS*) – see below, vinculin TFP control (*VinTFP*), vinculin tailless control (*VinTL*), *Wnt5a*^+/-53^, *Piezo1*^+/-95^. All mouse lines were outbred to CD1. All animal experiments were performed in accordance with protocols approved by the Animal Care Committ,ee Hospital for Sick Children.

#### Vinculin tension sensor knock-in mouse strains

To target the *Rosa26* locus, constructs were cloned using the following plasmids: vinculin tension sensor (*VinTS*) and vinculin tailless (*VinTL*)^68^ (Addgene plasmid#: 26019, 260020), and *Ai27* (gift from Hongkui Zeng^96^; Addgene plasmid#: 34630). To generate *VinTS* targeting vector, Mlu1 sites were introduced upstream of the start codon and downstream of the stop codon in *VinTS* using PCR. To generate *VinTL*, an Mlu1 site was introduced upstream of the start codon and a stop codon followed by an Mlu1 site downstream of the Venus sequence. To generate vinculin teal fluorescent protein (*VinTFP* – FRET donor only), an Mlu1 site was introduced upstream of the start codon and a stop codon followed by an Mlu1 site downstream of the mTFP1 sequence. PCR products were digested with Mlu1 and purified using Qiaex gel purification kit (Qiagen). Reverse primers for generating VinTL and *VinTFP* were identical, so the constructs were distinguished based on size in an agarose gel following Mlu1 digestion and gel purification and bands were cut out accordingly. The inserted construct in the *Ai27 Rosa26* targeting vector was removed with Mlu1 and replaced with *VinTS, VinTL*, or *VinTFP*. Sequences for all targeting vectors were confirmed through DNA sequencing (performed by The Centre for Applied Genomics, The Hospital for Sick Children). The final targeting vectors were constructed as follows: CAG enhancer – *FRT* – *loxP* – stop codons – 3x SV40 poly(A) – *loxP* – *VinTS/VinTL/VinTFP* – *WPRE* – *bGH* poly(A) – *attB* – *PGK* promoter – *FRT* – Neo – *PGK* poly(A) – *attP*.

Generation of ES cell lines and generation of chimeras were performed by the Transgenic Core at the Toronto Centre for Phenogenomics. Briefly, linearised constructs were electroporated into G4 ES cells and G418-resistent clones were screened by PCR. 4 positive clones from *VinTS*, and 2 positive clones from each of *VinTL* and *VinTFP* were aggregated with CD1 morula to obtain chimeric mice following standard procedures. Chimeric mice were outbred to CD1 mice to obtain an F1 generation through germline transmission. All procedures involving animals were performed in compliance with the Animals for Research Act of Ontario and the Guidelines of the Canadian Council on Animal Care. The Toronto Centre for Phenogenomics (TCP) Animal Care Committee and Hospital for Sick Children Animal Care Committee reviewed and approved all procedures conducted on animals at TCP and at the Hospital for Sick Children, respectively.

#### *Z/Wnt5a* conditional overexpression mouse strain

The *Z/Wnt5a* line was generated by targeting a *Cre*-activated *Wnt5a* expression vector to the endogenous *Ubiquitin-b (Ubb*) locus (unpublished reagent provided by Phil Smallwood and Jeremy Nathans). The *Cre*-activated *Wnt5a* expression vector was constructed using a similar strategy as described for generating the *Z/Norrin* expression cassette^97^. Expression of the *Wnt5a* transgene is activated only after *Cre-* mediated recombination, which removes a *LacZ-STOP* cassette upstream of the *Wnt5a* open reading frame. *Sox2:Cre*^98^ was employed to drive ubiquitous expression.

### Optical projection tomography

E9.5 mouse embryos were harvested and fixed in 4% paraformaldehyde overnight at 4°C. OPT was performed using a system that was custom-built and is fully described elsewhere^99^. Three-dimensional (3D) data sets were reconstructed from autofluorescence projection images acquired over a 25 minute scan period at an isotropic voxel size of 4.5 μm. The 3D surface renderings of the OPT data were generated by MATLAB software, version R2011b (Mathworks).

### Cell cycle time measurement

Pregnant females were injected first with BrdU intraperitoneally at E9.75 and then with IddU after 2.5hr. Thirty min. following the second injection, embryos were dissected in cold PBS and fixed with 4% PFA overnight at 4 °C. Whole-mount immunofluorescence of BrdU and IddU was performed^54^.

### Elasticity and viscosity measurement by atomic force microscopy

Mouse embryos were incubated in 50% rat serum in DMEM on a 35 mm dish in which 2% agarose was poured around the perimeter. The mandibular arch was immobilised to the agarose with pulled glass needles pinned through the flank adjacent to mandibular arch. The arch was examined using an AFM (BioScope Catalyst, Bruker) mounted on an inverted microscope (Nikon Eclipse-Ti). AFM indentation tests were performed using a spherical tip (radius: 15 μm) at distinct locations categorised as proximal, middle and distal mandibular arch. Spherical tips were made by assembling a borosilicate glass microsphere onto a tipless AFM cantilever using epoxy glue. The cantilever spring constant was calibrated before every experiment by measuring power spectral density of thermal noise fluctuation of the unloaded cantilever.

To determine the elastic modulus of epithelium, a trigger force of 300 pN was consistently applied. For both small and large indentation measurements in each region (proximal, middle, and distal), 15 locations were measured and repeated in triplicate, amounting to 45 measurements at each region. Because the epithelial thickness is approximately 10 μm, we followed the empirical 10% rule^100^ to indent epithelium up to 1 μm in depth to avoid influence from the mesenchyme. The Hertz model for a spherical tip was used to calculate the elastic modulus of the epithelium from the small indentation data. To determine mesenchymal elastic modulus and the overall tissue viscosity, we applied large indentation (depth: 7-10 μm) at indentation rates of 5, 10 and 15 μm/s. The Kelvin-Voigt model was used to fit the large indentation force-displacement data. To extract overall tissue viscosity, the range of data beyond 1.5 μm indentation depth was used, and the elastic range was neglected. The elastic modulus value of mesenchyme was then determined by using experimental force-displacement data and finite element simulation.

#### a) Determination of epithelium’s elastic modulus

For small indentation, the Hertz model for a spherical tip was used to fit the experimental force-displacement data and determine epithelial elastic modulus values. The relationship between the indentation depth *d* and the loading force *F* is

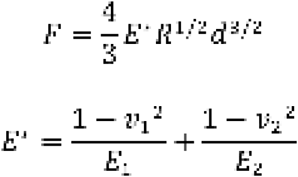

where *E_1_, v_1_* are the elastic modulus and Poisson’s ratio of the indenter; and *E_2_, v_2_*, are the elastic modulus and Poisson’s ratio of epithelium. The spherical tip was made of borosilicate glass (elastic modulus and Poisson’s ratio: 63 Gpa and 0.2). The Possion’s ratio for epithelium was set to 0.4^101^. Using the experimental data from small indentation as well as the above calculation, epithelial elastic modulus *E_2_* was quantified.

#### b) Determination of the overall embryonic tissue viscosity

For large indentation, the Kelvin-Voigt model was used to fit experimental force-displacement data. Stress-strain relation in the Kelvin-Voigt model is

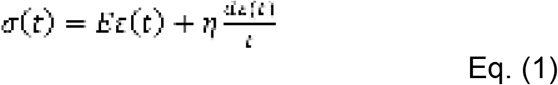

When the equation is multiplied by contact area *S = πa^2^*, where a is the contact radius, the left-hand side results in total force applied to epithelium and mesenchyme. For a bilayer structure, the contact radius 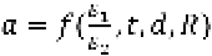 is a function of the first-second layer elastic mismatch ratio *E_1_/E_2_*, the first layer thickness *t*, indentation depth *d*, indenter radius R. Comparing the elastic modulus of epithelium *E*_epithelium_ and the overall modulus (i.e., epithelium and mesenchyme both included), we found the ratio *E*_epithelium_/*E*_mesenchyme_) was small (~2). Thus, according to the Hertz model^102^, 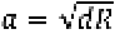. Rewriting Eq. (1) gives

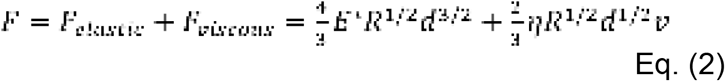

where *v* is indentation rate. It is evident that *E_elastic_* is rate-independent while *F_viscous_* is rate-dependent. Therefore, *F_elastic_* and *F_viscous_* were determined with the force-displacement data measured at different indentation rates, and viscosity η was also calculated. Furthermore, *F_elastic_* was exported into the finite element model to calculate mesenchyme’s elastic modulus.

#### c) Determination of mesenchyme’s elastic modulus

A snapshot of the meshed 2D axisymmetric finite element model with a bilayer structure (epithelium and mesenchyme) under frictionless loading is shown in Supplementary Fig. 1B. Supplementary Fig. 1F shows the steps we used to conduct iterative FE simulation to quantify mesenchyme’s elastic modulus. As in the above section, for each displacement, *F_elastic_* was determined. This elastic force applied to the overall tissue was incorporated into the finite element model. Since epithelial elastic modulus has been determined, different modulus values were assigned to the mesenchymal layer in the finite element model until finite element-obtained force-displacement curves agreed well with the experimental curves (Fig. 1F). Mesenchymal elastic modulus was saved once the R-square value (correlation coefficient between the two curves) was greater than the threshold value (0.99).

### Finite element modelling

An OPT (optical projection tomography) image stack was first concatenated and reconstructed using ImageJ, and a 3D model was exported in the STL format. We used MeshLab to isolate the mandibular arch from the embryo. Quadric edge collapse decimation algorithm was employed to reduce the total number of 3D triangular mesh faces to the target number of faces (~5,000 without distorting the geometry). The model was then imported into Solidworks in which the 3D mandibular arch model was segmented into two layers (epithelium and mesenchyme) with 24 regions based on the geometric locations of cell cycle time measurements. The finite element model was fixed in all six degrees of freedom (Displacement: *U_x_*=*U_y_*=*U_z_*=0, Rotation: *U_Rx_*=*U_Ry_*=*U_Rz_*=0) at the extended proximal end to ensure the boundary condition did not influence growth of the proximal region (Supplementary Figure 1G.). A ten-node tetrahedral element (SOLID 186) was selected to discretise the model.

Each region of the model was assumed to be viscoelastic, isotropic and homogeneous. The viscoelastic property of each region was assigned with an average value obtained from experimental AFM measurement and implemented in ANSYS v15.0 (ANSYS Inc., Canonsburg, PA) with instantaneous elasticity and two-pair Prony relaxation. We also performed comparative simulation with different viscoelastic values within the range of our AFM measurement and observed no significant effect in the predicted tissue shape change.

The experimentally measured 3D cell cycle time values were converted to strain using the method described previously^30^, and strain values were incorporated in the finite element model for tissue shape prediction. Cell density was considered to be the number of cells per volume, 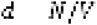 Therefore, tissue volume change was a function of cell number change and cell density change. We started by assuming cell density remains constant (changes in cell density were incorporated later as a correction factor), and tissue volume change was proportional to cell number change 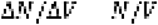 Volume strain 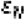 induced by cell number change was thus equal to 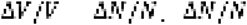 was calculated from cell doubling time. Cell density change (measured to be 1.7 % per hour) was incorporated into volume strain as a correction factor ε_d_. Hence, the overall volume strain ε was 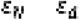.

Parameters incorporated into the model

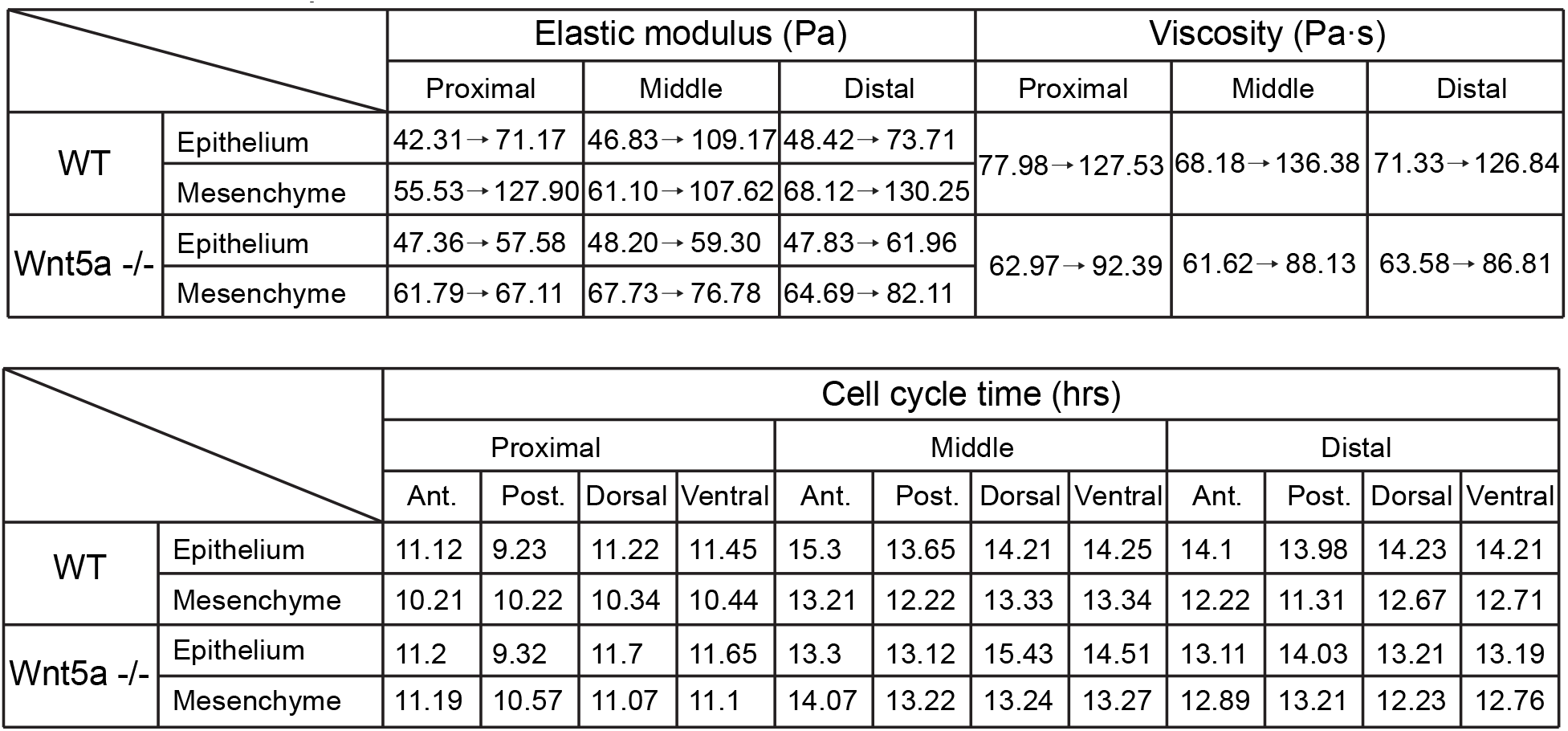

#### Voronoi tessellation and rigidity analysis

Using light sheet microscopy, we obtained approximately 1,000 2-dimensional images (ML-PD plane) of the mandibular arch, each image about 0.4 microns apart from the next along the perpendicular RC axis. Cell nuclei marked by *H2B-GFP* emitted light with relatively high intensity. Each image was processed as following: a) using a Gaussian deconvolution with a standard deviation roughly the size of one cell, the images were smoothened. b) Local maxima were sought in square boxes, again, roughly the size of one cell. Each local maximum was marked as a cell centre. This process was repeated for each image in ML-PD plane. The layers of marked cells were then compared against one other. If a point was marked to be the centre of a cell within 7 or more consecutive layers in the stack, the middle layer was marked as the RC position of that cell. This process gave us a 3D representation of roughly 6,000 cell nuclei in the tissue. Voronoi tessellation was performed using these cell nuclei as nodes for the tessellation algorithm.

#### Live, time-lapse confocal microscopy

Live image acquisition was performed as described previously^14,32^. Briefly, embryos were submerged in 50% rat serum in DMEM without phenol red (Invitrogen) in a 25 mm imaging chamber. Cheese cloth was used to immobilise the embryo and position the mandibular arch directly against the coverglass. Embryos were imaged in a humidified chamber at 37°C in 5% CO_2_. Time-lapse images were acquired on a Quorum WaveFX-X1 spinning disk confocal system (Quorum Technologies Inc.) at 20X magnification. Images were processed with Volocity software or ImageJ/Fiji. Representative images are shown from at least 3 independent experiments for each condition, and unless otherwise indicated, from at least 3 independent cohorts. No statistical method was used to predetermine sample size. Experiments were not randomised. Investigators were not blinded to allocation during experiments and outcome assessment.

#### Live, time-lapse light sheet microscopy

Three-dimensional (3D) time-lapse microscopy was performed on a Zeiss Lightsheet Z.1. microscope. Embryos were suspended in a solution of DMEM without phenol red containing 12.5% rat serum and 1% low-melt agarose (Invitrogen) in a glass capillary tube. Once the agarose had solidified, the capillary was submerged into an imaging chamber containing DMEM without phenol red, and the agarose plug was partially extruded from the capillary until the portion containing the embryo was completely outside of the capillary. The temperature of the imaging chamber was maintained at 37° C with 5% CO_2_. Images were acquired using a 20X/1.0 objective with dual-side illumination, and a z-interval of 0.479 μm. All experiments were imaged in multi-view mode with 3 evenly-spaced views spanning approximately 90 degrees (from a frontal view to a sagittal view of the mandibular arch). Images were acquired for 3-4 hours with 10 minute intervals. Fluorescent beads (Fluospheres 1μm, Thermofisher, 1:10^6^) were used as fiducial markers for 3D reconstruction and to aid in drift-correction for cell tracking. Multi-view processing was performed with Zen 2014 SP1 software to merge the 3 separate views and generate a single 3-dimensional image. Further analysis and cell tracking were performed using Arivis Vision4D software (Arivis).

#### Membrane segmentation and 3D cell neighbour counting

3D timelapse datasets of cell membranes were processed with the ImageJ macro ‘TissueCellSegmentMovie’ (kindly provided by Dr. Sébastien Tosi from the Advanced Digital Microscopy Core Facility of the IRB Barcelona) to generate membrane segments prior to analysis with Imaris software (Bitplane). Surface objects were created using Imaris and this data was used for 3D cell neighbour analysis. Cell neighbours were manually counted by identfying the target cell and counting all cells which share an interface within the plane, as well as adjacent cells in the planes above and below. Analysis was performed on 3 separate areas in both middle and distal regions of the mandibular arch over 2 independent experiments for each condition.

#### Cell tracking and dandelion plots

Cell tracks (cell positions tracked over time) were calculated manually. Cell nuclei were followed between z-stacks of images as described above, each z-stack separated from the next by 5 or 10 minutes. The large drift between images over time and the resolution of the images made it difficult to follow more than 130 cells throughout the entire movie. Only cells that could reliably be identified and followed by eye in at least 33 frames were considered for the random walk analysis.

For each movie, the z-stacks for each time point were aligned, and ‘fusion’ stacks were created to generate z-plane images based on the mean voxel intensity at each voxel. The ‘fused’ stacks were then imported into Arivis Vision 4D software (arivis AG, Unterschleißheim, Germany), and automated tracking was used to generate cell tracks. Tracks were validated manually, and validity annotations were entered into an Excel spreadsheet. Tracks were defined to be valid at a given timepoint if the segment determining the track’s location at that timepoint was centred in the middle of a nucleus. Tracks were defined to be suitable for use in further analysis if time between their first valid timepoint and final valid timepoint was 150 minutes or greater. Whether a track tracked an epithelial or mesenchymal cell was determined observing the nuclear location of the track relative to the tissue boundary as well as the morphology of the nucleus (epithelial cells tended to have more elongated, columnar shapes relative to their counterparts in the mesenchyme).

To correct for drift, the frame by frame displacement of red fluorescent beads that were embedded adjacent to the embryo within the agarose plug used in the light sheet microscope was calculated. These displacements were imported into MATLAB for initial track concatenation and drift correction. Track and segment information and validity were acquired on Arivis. The concatenation function returned drift corrected tracks that were filtered to only include valid tracks. Tracks could then be plotted, and end to end displacements calculated and plotted.

#### Random walk model

We described cell motions from tracking experiments by stochastic processes. In particular, we adopted the Ornstein-Uhlenbeck (OU) process^103^ for our analysis. In an OU-process, the trajectory of a particle is determined by a relation describing its change in velocity over time. The change of velocity in the model is proportional to its velocity in the past (persistent) plus a randomly generated term (random walk). This approach has been used to analyse the properties of cell migrations under many contexts including tumour growth^62^ and wound rep ait^61^.

The OU-process is defined by the Langevin equation

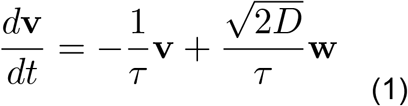

where v is the cell velocity, t is time, D is the diffusion coefficient, vector w describes a Wiener process, and 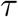 is the time scale often referred to as the persistent time. The persistent time, which may be understood as the length of time a given velocity “remembers” itself, describes the time of the velocity auto-correlation function

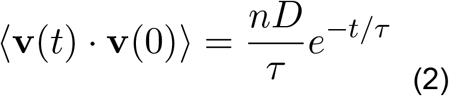

where n is the space dimension of the tracks. The mean squared displacement (MSD) is given by

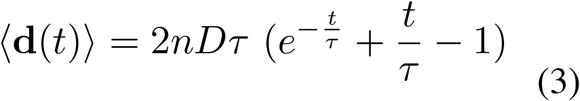

Equation (2) and (3) was used to fit the observed cell tracks to obtain the persistent time 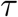 and the diffusivity coefficient D. Consequently, we were able to simulate cell trajectories by equation (1) to further understnnd similarities and discrepancies between the statistics from the observations and the OU-process.

The wild type dataset that we used for the numerical simulation contained n = 179 number of cells tracked with a time interval of t = 5 min. over a 3 h period. Following Wu et al., 2014^62^, we simulated cell trajectories by applying a first-order Euler’s scheme to equation (1) to obtain

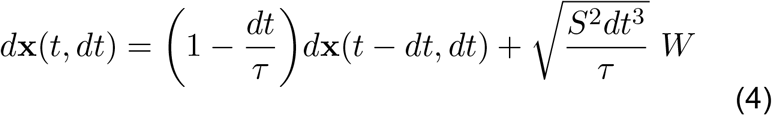

where W ~ *N*(0,1) and *S* is the cell speed with 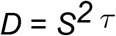. We applied principle component analysis to observed cell trajectories prior to our analysis so that we could simulate cell displacement in each direction separately. In other words, we diagonalised the correlation coefficiant matrix so that off-diagonal components were zero.

An important charcteristic of a random walk model is its mean squared distance (MSD). For an arbitrary trajectory, the MSD as a function of time follows a power-law, with power of 0 representing a truly random trajectory, and power of 2 representing movement in a straight line with no randomness. On a log-log plot, these two extremes translate into lines with slope 0, or a horizontal line, for the random trajectory, and a line of slope 2 for the walk in a straight line. A persistent random walk, however, is characterized by slope 1, the short black line in the figure. The analysis of cell trajectories (based on light sheet microscopy) suggested that mandibular arch cells exhibited a MSD slope that is characteristic of a persistent random walk.

#### Strain analysis

A rectangular grid of points was superimposed on the first frame of a rostrocaudal-proximodistal plane of a time-lapse light sheet movie of the mandibular arch. The points were followed in subsequent frames by calculating the correlation function of a box around each point in the initial time frame to all the boxes of the same size in the vicinity of the original box in the next frame. The vector that connected the location of the box in the first frame to the location of the neighbouring box with the highest correlation with the original box in the second frame was the displacement vector. The points were moved by this displacement vector in each time frame, and this process was repeated for all 31 time frames.

#### Immunostaining

Embryonic day (E) 9.0-9.5 mouse embryos were fixed overnight in 4% paraformaldehyde in PBS followed by 3 washes in PBS. Embryos were permeabilised in 0.1% Triton X-100 in PBS for 20 min and blocked in 5% normal donkey serum (in 0.05% Triton X-100 in PBS) for 1h. Embryos were incubated in primary antibody overnight incubation at 4°C. Embryos were washed in 0.05% Triton X-100 in PBS (4 washes, 20 min each), and then incubated in secondary antibody (1:1000) for 1 h at room temperature. Embryos were washed (4 washes, 20 min each), followed by a final wash overnight at 4°C, and stored in PBS. Images were acquired using a Quorum spinning Disk confocal microscope (Zeiss) at 10 X, 20 X or 40 X magnification, and image analysis was performed using Volocity software (Perkin Elmer) and ImageJ.

#### Antibodies

Phospho-myosin light chain 2 (Thr18/Ser19) (Cell Signaling #3671, rabbit, 1:250); PIEZO1 (FAM38A, proteintech #15939-1-AP, rabbit, 1:250); N-cadherin (BD #519001943, mouse 1:250); Desmoglein (BD #51-9001952, mouse, 1:250); BrdU (B44) (BD #347580, mouse, 1:250); Calpain2 (E-10) (Santa Cruz #sc-373966, mouse, 1:50); VANGL2 (Sigma HPA027043, rabbit, 1:250). All secondary antibodies were purchased from Jackson Immunoresearch and used at 1:1,000 dilutions. Rhodamine phalloidin (Invitrogen #R415, 1:1,000); Alexa Fluor 633 phalloidin (Invitrogen #A22284, 1:1000).

#### Whole mount in situ hybridisation

Whole mount in situ hybridisation was performed as described^104^. Wildtype and mutant littermate embryos were processed identically in the same assay for comparison. The *Wnt5a* riboprobe was gift from A. P. McMahon.

#### Quantification of polarised actin and myosin distribution

Single confocal slices taken 2 μm above (for epithelium) and below (for mesenchyme) the basal surface of epithelial cells stained with phospho-myosin light chain 2 (Thr18/Ser19) (pMLC), rhodamine-phalloidin or Alexa Fluor 633 phalloidin were analysed using SIESTA software as described previously^14,105^. Cell interfaces were manually identified, and average fluorescence intensities were calculated for all interfaces and grouped into 15° angular bins, with 0°-15° bin representing interfaces that are parallel with the AP axis (PD interfaces) and 75°-90° bin representing interfaces that are parallel with the PD axis (AP interfaces). Average fluorescence intensity values for each bin were normalised to average fluorescence intensity of PD interfaces (0°-15° angular bin). Error bars indicate standard error of the mean and p values were calculated using Student’s t-test.

#### Quantification of cell behaviours

Metaphase-to-telophase transition angles were measured as described^32^. Epithelial tetrad formation and resolution angles were measured by first assigning a proximodistal reference axis taken from a low 10 X confocal magnification view of the embryo flank. Tetrads were identified manually, frame by frame. The angle between the long axis of the ellipse outlined by each tetrad and the reference axis was documented at the beginning of a given movie and upon resolution.

#### Live, time-lapse fluorescence lifetime analyses *in vitro* and *in vivo*

Fluorescence lifetime microscopy (FLIM) was performed on a Nikon A1R Si laser scanning confocal microscope equipped with PicoHarp 300 TCSPC module and a 440 nm pulsed diode laser (Picoquant). Data were acquired with a 40 x/1.25 water immersion objective with a pixel dwell time of 12.1 μs/pixel, 512 x 512 resolution, and a repetition rate of 20 MHz. Fluorescence emission of mTFP1 was collected through a 482/35 bandpass filter. Embryos were prepared as described above for live timelapse microscopy and imaged in a humidified chamber at 37°C in 5% CO_2_. Data were acquired over 15 frames at 2 minute intervals for 20 minutes.

Fluorescence lifetime of mTFP1 was determined using n-exponential reconvolution model in SymphoTime software (Picoquant) with model parameters n=2 for VinTS and VinTL and n=1 for VinTFP. Fitting was performed to achieve a chi-squared of χ^2^=1000 ± 0.100.

FRET efficiency (E) was calculated based on the molecular separation between donor and acceptor and the corresponding fluorescence lifetime using the following equation:

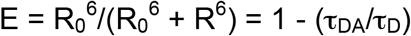

where R_0_ = Förster radius of donor-acceptor pair, R = donor to acceptor separation distance, TDA = fluorescence lifetime of donor in the presence of acceptor, and TD = fluorescence lifetime of donor only.

Force was calculated from FRET efficiency based on the calibration results reported by Grashoff et al ^68^.

#### Inhibitor treatment

The effect of inhibitors Y27632 (Sigma #Y0503) and Cytochalasin D on fluorescence lifetime of VinTS were conducted as follows: Embryos were incubated with 20 μM Y27632 or 20 μg/mL Cytochalasin D for 15 minutes in 50% rat serum with DMEM in roller culture prior to experiments that were conducted using the same conditions described above.

For Calpain inhibition, embryos were treated in roller culture with 10 mM PD 150606 (Calbiochem #513022), 10 mM PD151746 (Abcam #ab145523), 10 mM MDL28710 (calpain inhibitor III, Sigma #208722) or DMSO control in culture medium for 3 h, then fixed in 4% paraformaldehyde overnight at 4° C.

#### Cytosolic calcium fluctuation

Calcium indicators Fluo-8 AM (Abcam #142773) or X-rhod-1 AM (Invitrogen #X14210) at 10 mM were added in 2 ml of 50% rat serum medium and incubated for 30 min. at 37 °C. Embryos was permiabilised with 0.1% Pluronic F-127 (Invitrogen #P3000MP) in culture medium for 20 min. before X-rhod-1 staining. The culture medium was removed and washed (2 times) and replaced with 2 ml culture medium. Live cell calcium imaging was performed on a Quorum Spinning Disk confocal microscope (Zeiss) equipped with a 40 X water objective lens or a Nikon A1 confocal microscope (Nikon) equipped with a X dry objective lens. Calcium indicators were excited with an argon laser line (488 nm), and emissions were recorded in the green channel (500-560 nm) for Fluo-8 AM (acetoxyomethyl) and red channel (600-700 nm) for X-rhod-1 AM. Images were acquired every 2 or 5 min. for up to 20 min.; image and data acquisition was performed using Volocity software (Perkin Elmer). All imaging experiments were performed in duplicate.

#### VinTS/Ca^2+^ correlation

We calculated the correlation between VinTS lifetime and Ca^2+^ concentration data series with zero time-lag. In this study, we defined the percentage of X-rhod-1 AM staining area for each cell in relation to Ca^2+^ concentration. Assume the first time series (force vs. time) are called x (x1, x2, x3, etc. for time points 1, 2, 3), and the second series are y (Ca^2+^ concentration vs time). First we calculated the mean and standard deviation of x and y. Let us refer to the mean of x, μ_x_ and standard deviation of x as σ_x_. μ_y_ and σ_y_ for y. N is the number of data points. We subtracted the means from each series and multiplied data points corresponding to the same time, and summed the results. The formula follows:

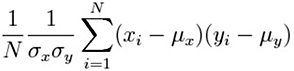

We analysed the amplitude of normalised correlation by using MATLAB xcorr (https://www.mathworks.com/help/signal/ref/xcorr.html).

This will give us a number between −1 and 1. For values approaching 1, the two series are strongly correlated. Approaching 0, they are not correlated. Approaching −1 implies they are anti-correlated.

---

**Supplementary Figure 1.**
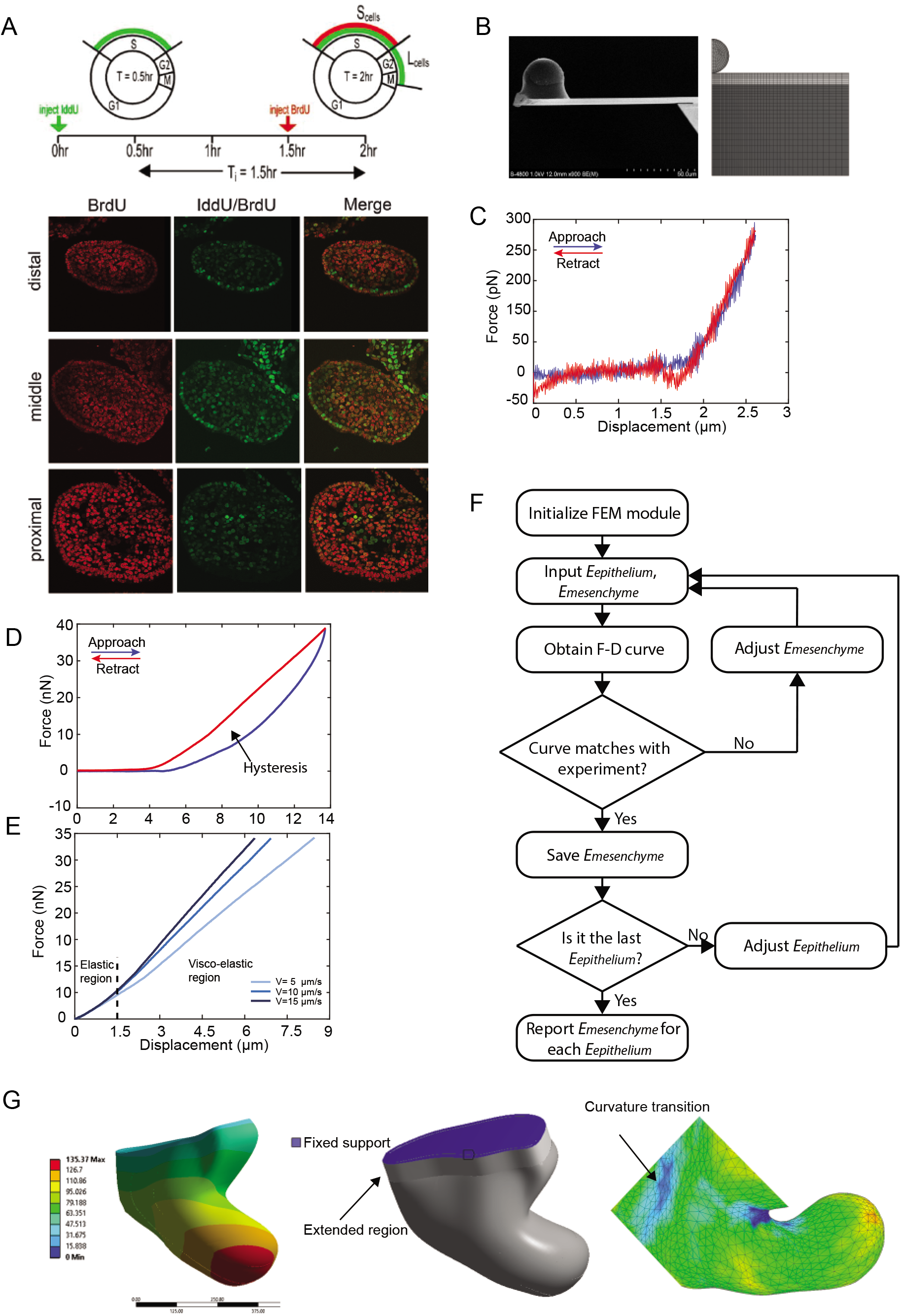
Viscoelastic properties of the mandibular arch. **A** Method of cell cycle time measurement using dual thymidine analog labelling to mark cells in S phase. The proportion of Iddu– and Brdu-positive cells were quantified after immunostaining to calculate cell cycle time in different regions of the mandibular arch. **B** Scanning electron micrograph of an AFM cantilever with a 30 μm spherical tip employed for embryo indentation (left) and the axisymmetric mesh of the finite element model (right). **C** A representative force-displacement (F-D) curve of small indentation is consistent with purely elastic behaviour. **D** A representative force-displacement curve under large indentation reveals evidence of hysteresis, or energy dissipation that indicates viscoelastic behaviour. **E** Force-displacement curves at different indentation rates indicate viscoelastic behaviour. **F** Flowchart of finite element model (FEM) simulation for determining the elastic modulus of mesenchyme. **G** Finite element-predicted deformation (in μm) (left); boundary condition set for simulation (centre); curvature transition used for determining the proximal end (right).

**Supplementary Figure 2.**
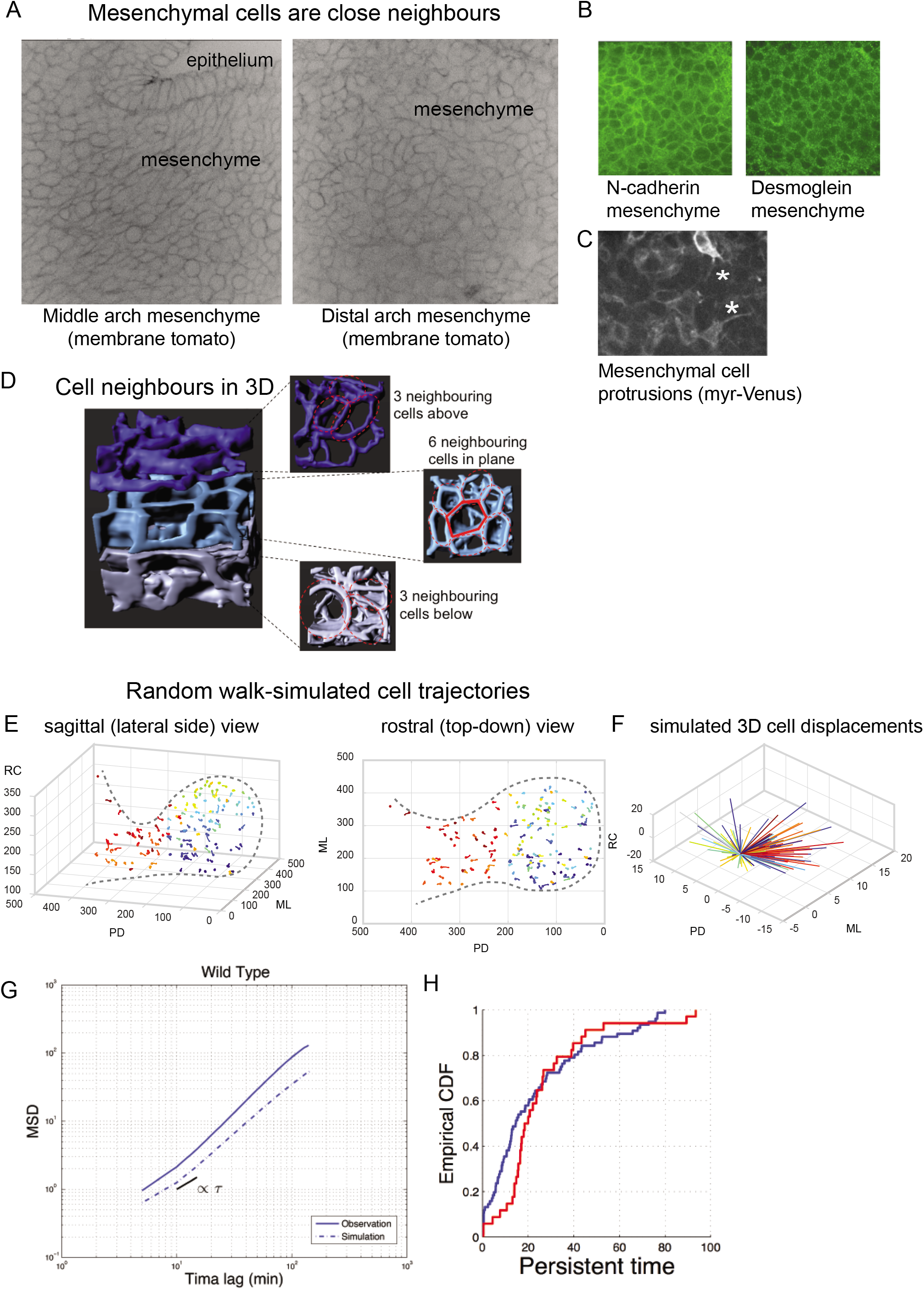
Mesenchymal cell neighbour relationships and random walk model. **A** Mesenchymal cells in live *mTmG* embryos are intimate neighbours without visible intercellular spaces by confocal microscopy. **B** Mesenchymal cells express abundant cell-cell adhesion proteins N-cadherin (left) and desmoglein (right). **C** Mosaic expression of *myr-Venus* among mesenchymal cells in the arch revealed protrusive activity that may facilitate cell rearrangements. **D** Cell neighbours in 3D were counted by examining small volumes of membrane segmented mesenchymal tissue among live *mTmG* embryos. **E**, **F** Simulated cell trajectories based on our random walk model (compare with Fig. 2F, G). **G** According to a the model that we employed (methods), mean squared displacement (MSD), a measure of the extent of random motion, was similar between observed and simulated cell tranjectories, thereby validating this model for statistical comparison between different regions of the arch. **H** Cells in the middle waist region exhibited a greater slope for cumulative distribution function (CDF) of persistent time compared to those in the distal bulbous region (p<0.01), implying motion of waist cells is more consistent over time compared to those of the bulbous region.

**Supplementary Figure 3.**
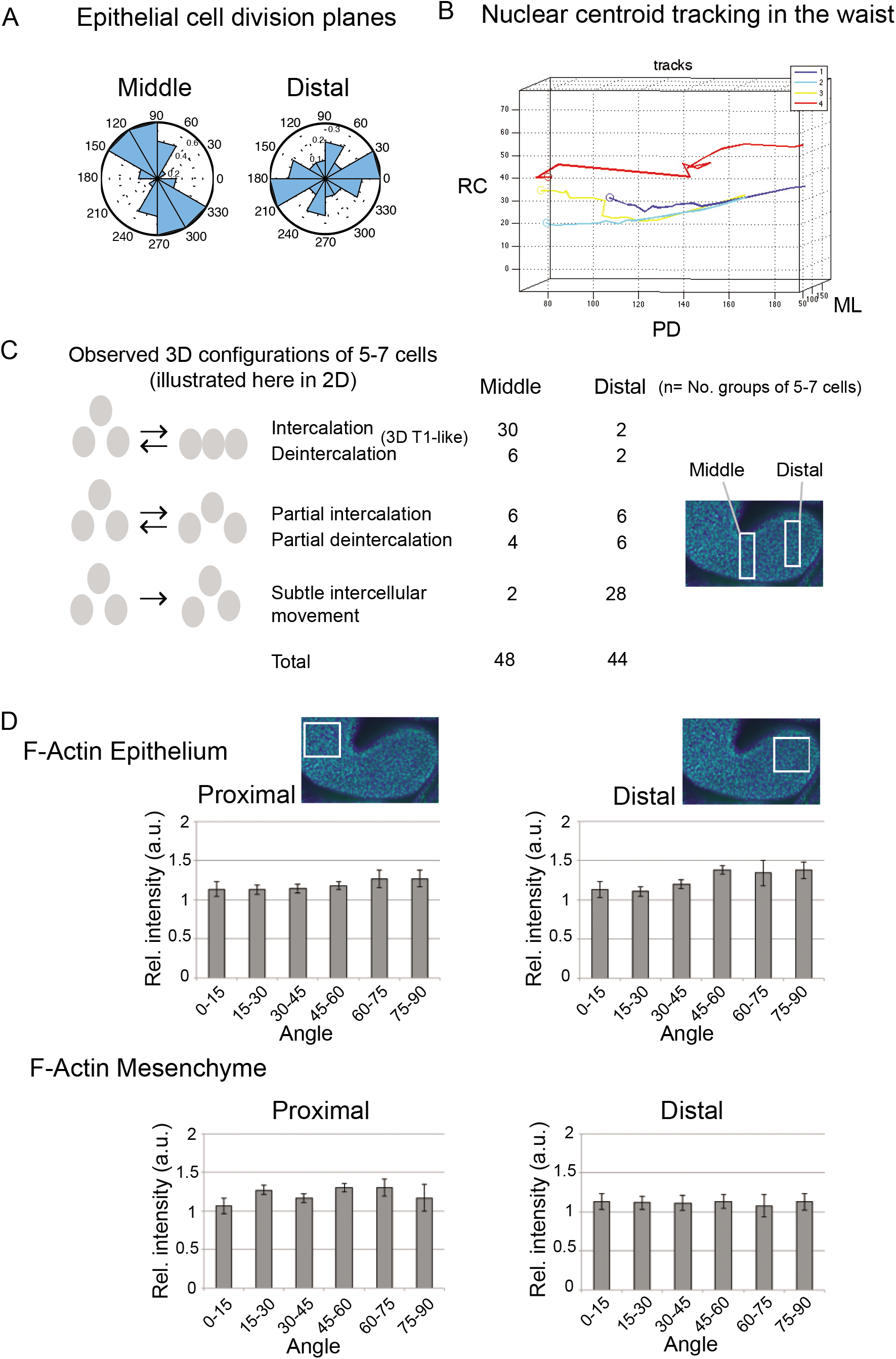
**A** Epithelial cell division planes in the middle and distal regions of the mandibular arch. **B** Tracks of neighbouring mesenchymal nuclei reveal AP convergence of cell tracks during proximodistal elongation of the middle region. **C** Proportion of different dynamic configurations of 5-7 mesenchymal cells that we defined in the middle and distal region. **D** The angular distribution of immunostain fluorescence intensity for epithelial (n=4 embryos) and mesenchymal (n=4 embryos) F-actin in proximal and distal regions of the arch relative to the arch long axis that was designated as 0° was quantified using SIESTA. Asterisks denote p<0.05, error bars denote s.e.m.

**Supplementary Figure 4.**
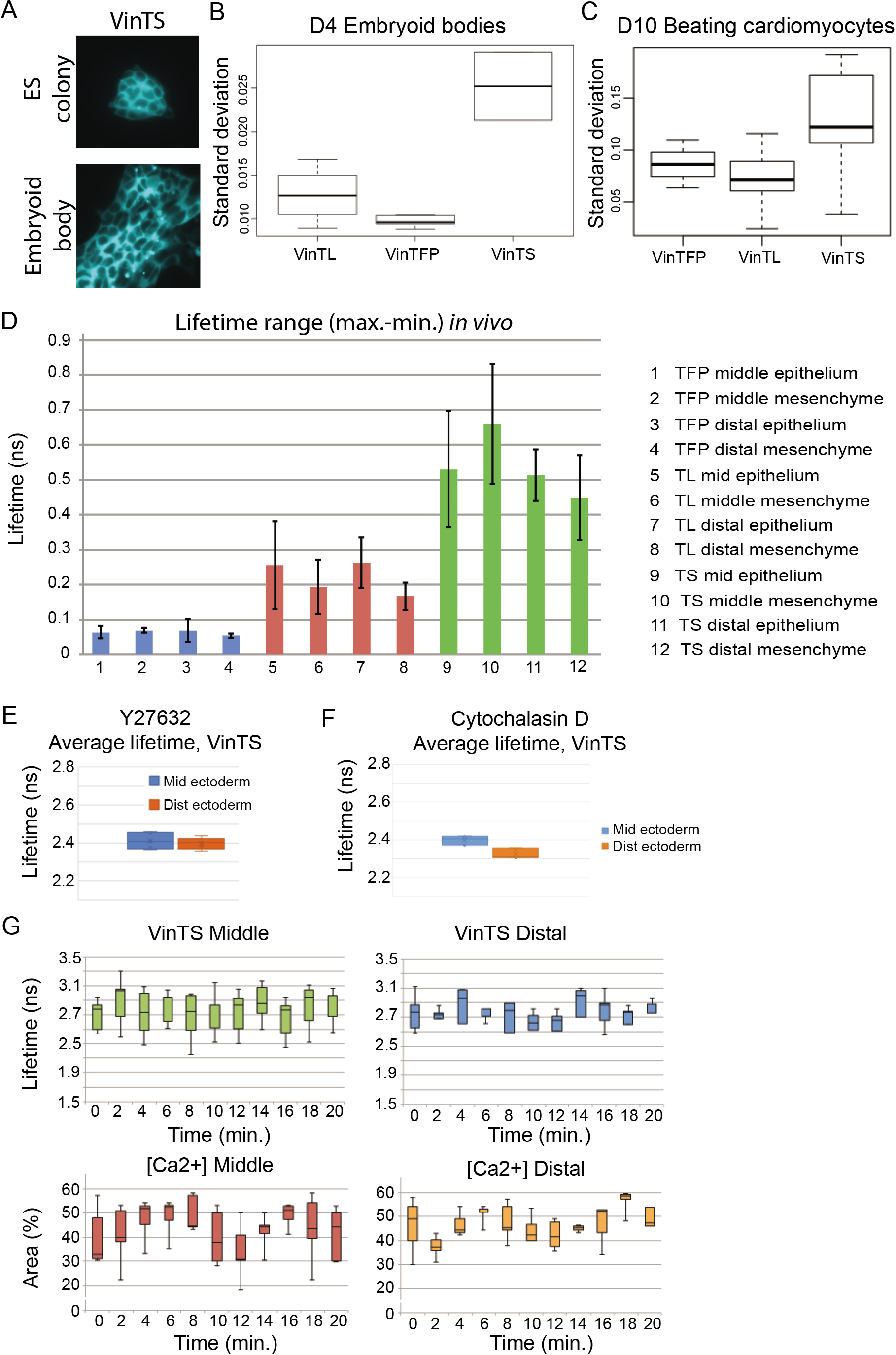
Vinculin force sensor evaluation in vitro and in vivo. **A** Expression of the full length tension sensor knock-in VinTS in an ES cell colony and an embryoid body. **B**, **C** Standard deviation of full length vinculin tension sensor (VinTS), TFP (FRET donor) only control (VinTFP), and vinculin tailless control (VinTL) in embryoid body cells (B) and differentiated beating cardiomyocytes (C). **D** Range of lifetimes values observed in middle (mid) and distal (dist) epithelium (ecto) and mesenchyme (meso) of the 20 somite stage mandibular arch for each of the VinTFP, VinTL and VinTS mouse lines, n=15 cells in each of 3 embryos, error bars indicate standard deviation. **E**, **F** VinTS lifetime in the mid-portion of the arch was dampened by treatment of embryos with Y27632, a ROCK inhibitor (E) and cytochalasin D, an actin polymerisation inhibitor (F) (n=15 cells per region in each of 3 embryos, error bars denote s.e.m; compare with 4C). **G** VinTS fluorescence lifetime and X-rhod-1 fluctuation data that correspond to correlations in Fig. 4F.

**Supplementary Figure 5.**
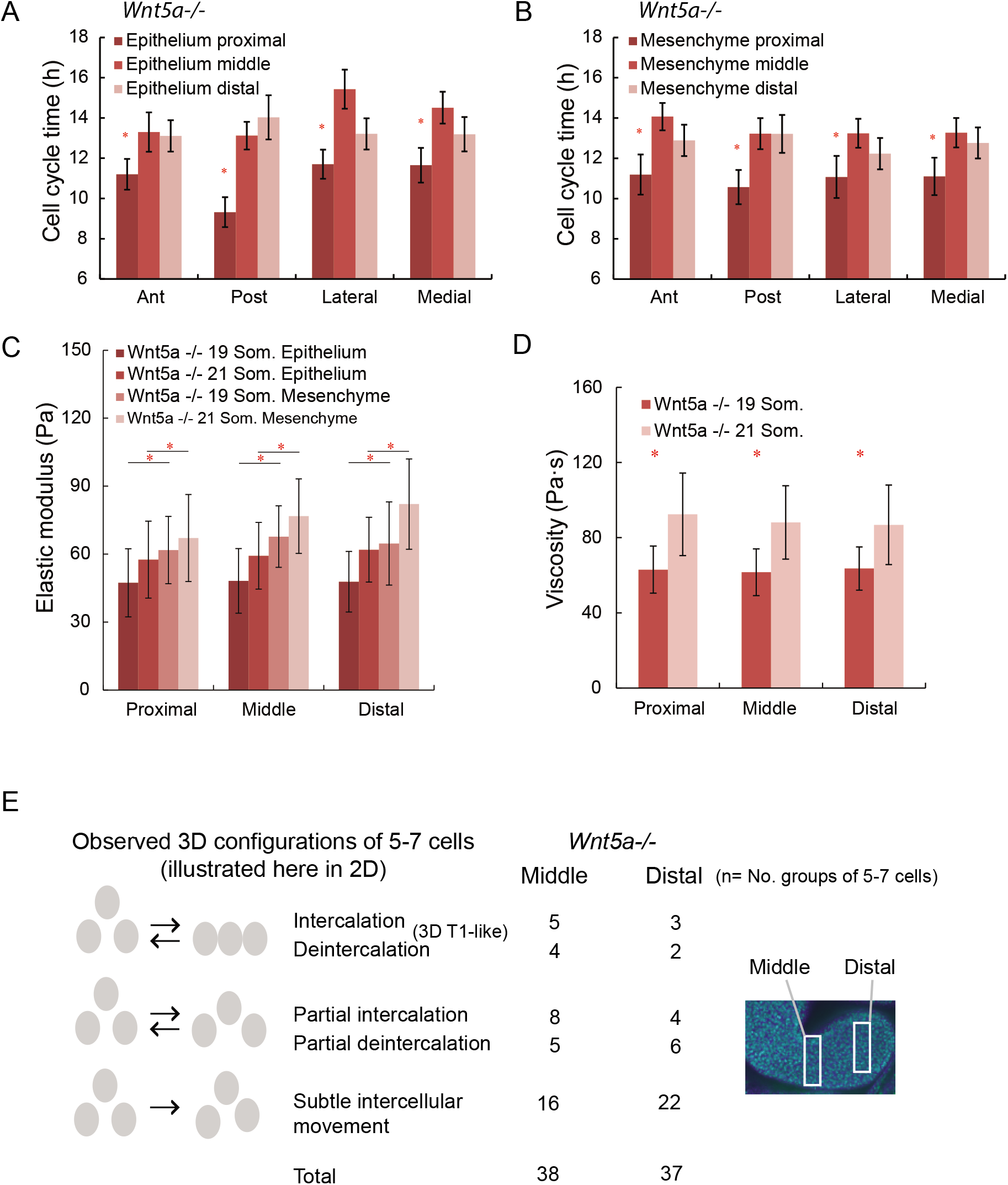
Spatial variation of cell cycle time, elastic modulus and viscosity in *Wnt5a^-/-^* mutant mandibular arch. **A**, **B** Epithelial (A) and mesenchymal (B) cell cycle times in in 24 adjacent regions of the mandibular arch (as in Fig. 1B). As in WT embryos (see Fig. 1C, D), cell division was more rapid in the proximal region for both epithelial and mesenchymal layers; n=3 embryos at 20 somite stage, 15-35 cells examined for each of 12 epithelial regions per embryo, 50-75 cells examined for each of 12 mesenchymal regions per embryo; asterisks denote p<0.05. **C** Elastic (Young’s) modulus (stiffness) of epithelium and mesenchyme. **D** Viscosity of whole tissue in proximal, middle and distal regions of the mandibular arch at 19 and 21 somite stages. In contrast to WT embryos (see Fig. 1F), middle arch epithelial stiffness did not increase significantly between 19 and 20 somite stages. For C and D, 15 separate sites in each proximal, middle and distal region were indented in triplicate (45 measurements per region) per embryo; n=3 embryos. Asterisks denote p<0.05, error bars denote standard deviation. **E** Proportion of dynamic configurations of 5-7 cells in the middle and distal regions of the *Wnt5a^-/-^* mutant mandibular arch.

**Supplementary Figure 6.**
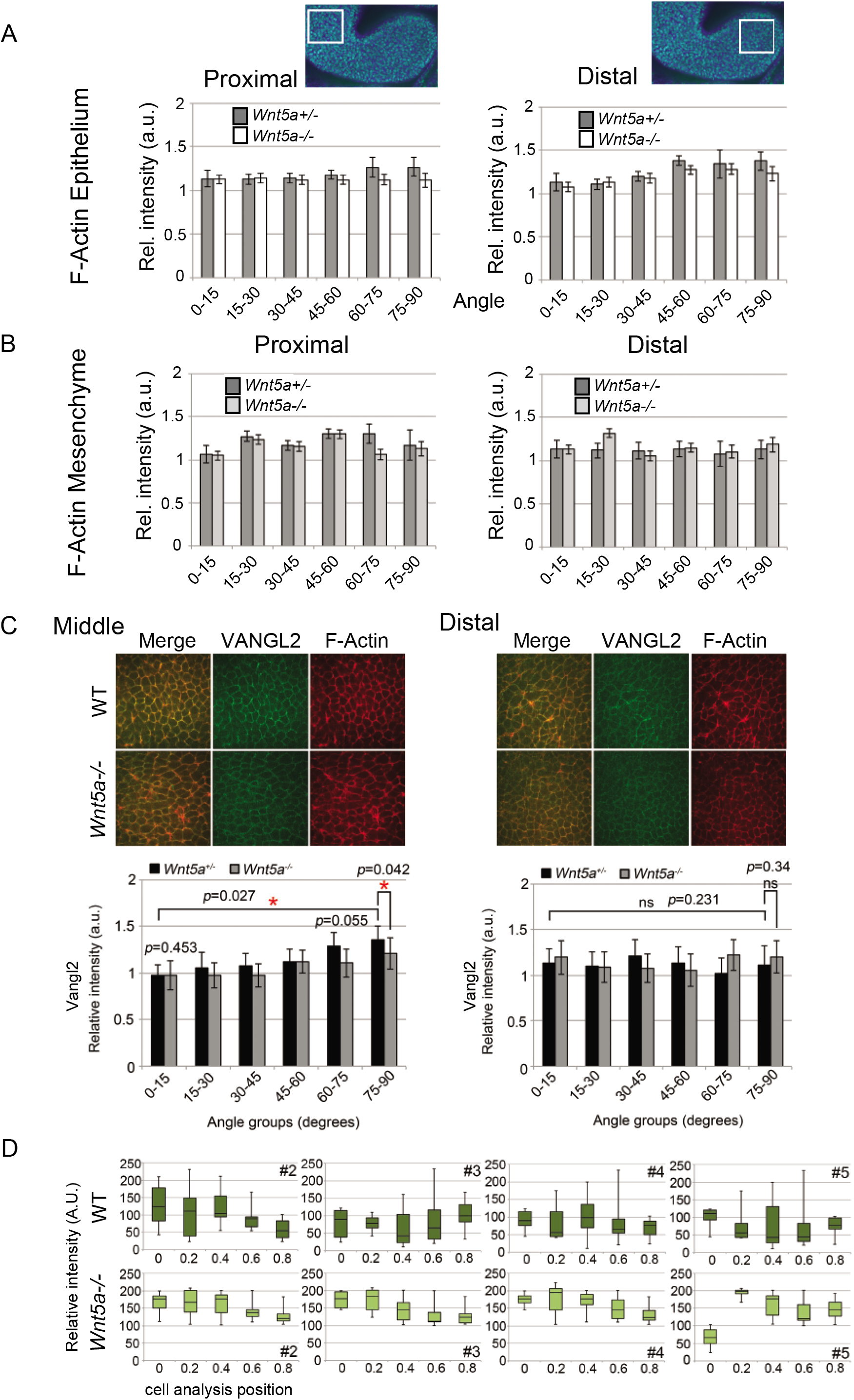
Cortical polarity is not biased proximal and distal regions of the mandibular arch. **A**, **B** F-actin orientation in proximal and distal regions for epithelium (A) and mesenchyme (B) in WT and *Wnt5a^-/-^* mutant embryos that corresponds to Fig. 6A. **C** VANGL2 immonostain intensity was biased parallel to actomyosin along proximodistal cell interfaces in the WT middle, but not distal, arch. That bias was diminished in *Wnt5a^-/-^* mutants. **D** Variation of cytosolic calcium concentration in individual cells over time; corresponds to Fig. 6E.

**Supplementary Figure 7.**
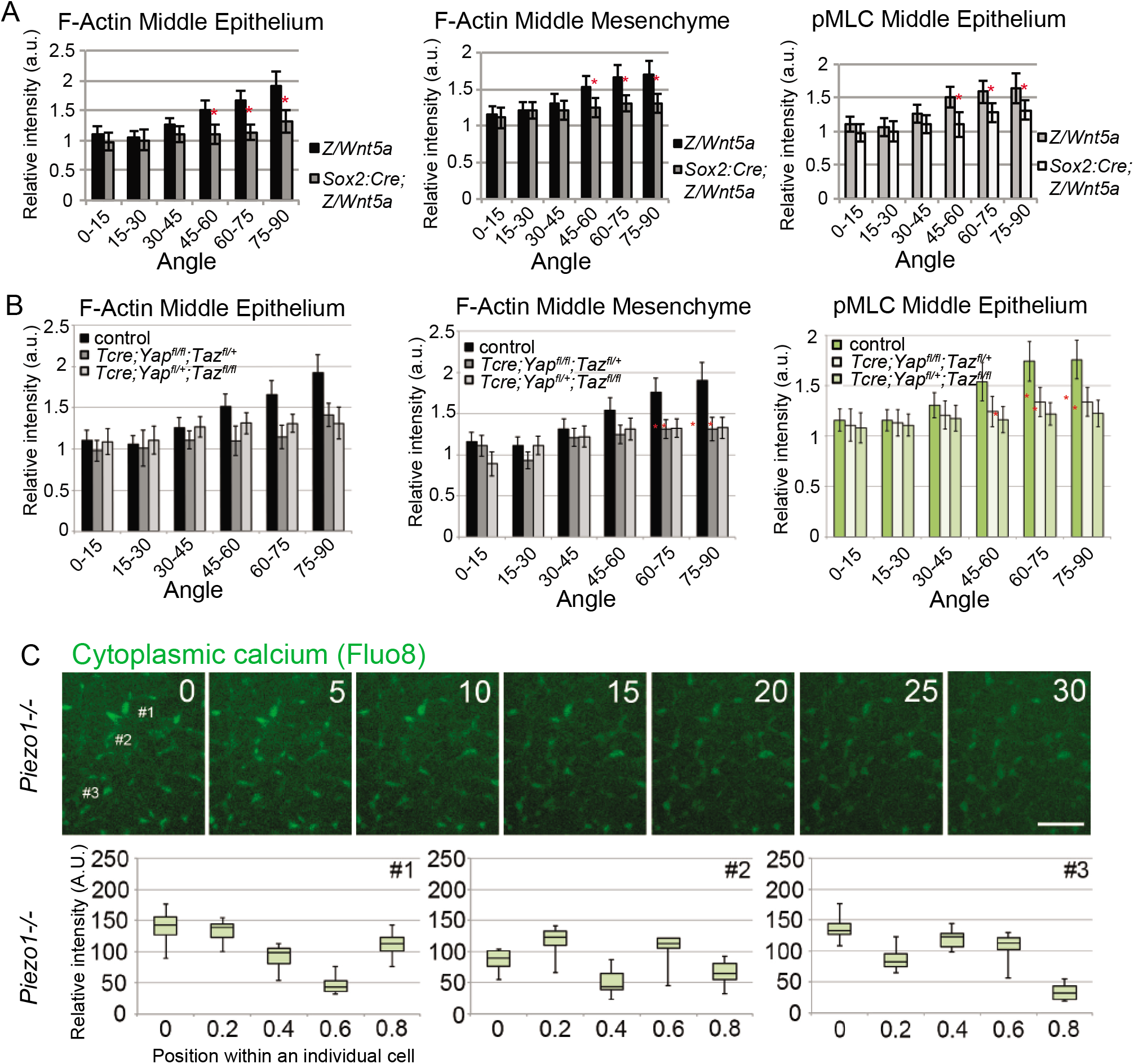
*Piezo1* regulates calcium flux. **A** F-actin and pMLC biases were diminished in 20-21 somite *Sox:Cre;Z/Wnt5a* embryos. **B** F-actin and pMLC biases were diminished in 20 somite *T:Cre;Yap^f/f^;Taz^f/+^* and *T:Cre;Yap^f/+^;Taz^f/f^* embryos. Asterisks indicate p<0.05. **C** Cytosolic calcium fluctuation in the 20-21 somite *Piezo1^-/-^* arch was dampened relative to WT embryos.

## MOVIE LEGENDS

**Movie 1** Rendering of 3D OPT image of the right mandibular arch of a 19 somite stage WT embryo.

**Movie 2** Rendering of 3D OPT image of the right mandibular arch of a 21 somite stage WT embryo.

**Movie 3** Rendering of 3D OPT image of the predicted shape of the right mandibular arch of a 21 somite embryo based on our finite element model. Four hours of growth (between 19-21 somite stages) were simulated starting with the shape of the actual WT 19 somite stage arch. Inputs included spatial distributions of cell cycle time, epithelial and mesenchymal Young’s modulus, and whole tissue viscosity.

**Movie 4** Three dimensional rendering of the right mandibular arch of a 20 somite *CAG::H2B-GFP* embryo imaged by live light sheet microscopy. Nuclei in the narrow mid-portion were oriented with their long axes perpendicular to the distalward axis of growth.

**Movie 5** Cell membranes have been segmented in this 3D rendering of the surface epithelium together with a partial volume of the underlying mesenchyme in a 20 somite embryo harbouring the *mTmG* transgene for membranes and *CAG::H2B-GFP* for nuclei. Time lapse light sheet microscopy of the intact embryo was performed. Distal is to the lower left.

**Movie 6** The neighbours of one mesenchymal cell (blue) have been coloured to facilitate counting.

**Movie 7** Three dimensional trajectories of a subset of nuclei labelled with H2B-GFP in a 20 somite WT embryo. Colour coding corresponds to plot in Mov. 8.

**Movie 8** Three dimensional start point-to-end point displacements (dandelion plot) of cells labelled in Mov. 7 with corresponding colour code. Red/orange cells representing the middle region move predominantly distalward whereas green/blue cells in the distal region move radially.

**Movie 9** Simulation of cell tracks that was based on our random walk model and was started from the initial frame of the same time lapse experiment shown in Mov. 7.

**Movie 10** Dandelion plot derived from random walk simulation shown in Mov. 9.

**Movie 11** Proximodistal-rostrocaudal plane taken from a 150 min. time-lapse, light sheet movie of the 20 somite stage mandibular arch. The superimposed strain grid is based on nodes that remain fixed to the same positions throughout the movie and highlight tissue deformation over time.

**Movie 12** An intermediate-scale volume of tissue of the middle arch of a 20 somite *CAG::H2B-GFP* embryo. Distal is to the right. The movie begins with an approximately rostral (top-down) view then turns to stabilise on a sagittal (lateral) view for the majority of the movie before returning back to a rostral view at the end.

**Movie 13** A small-scale volume of several cells from within the middle region of the mandibular arch of a 20 somite *CAG::H2B-GFP* embryo. The predominant view is sagittal, and distal is to the right. One central nucleus intercalated among several others in the direction of its long axis that is perpendicular to the axis of growth.

**Movie 14** Another example of a small-scale volume of several cells from within the middle region of the mandibular arch of a 20 somite *CAG::H2B-GFP* embryo. This movie rotates view about the rostrocaudal axis. One central nucleus intercalated among several others in the direction of its long axis that is perpendicular to the axis of growth.

**Movie 15** Combined H2B-GFP and mTmG labels with membrane rendering (red) shows one cell (arrow at the beginning) intercalate and separate neighbours caudal to (below) it within the middle region of the right mandibular arch of a 20 somite WT embryo. The view is sagittal-rostral oblique to best show the intercalation event.

**Movie 16** A small-scale volume of several cells from within the distal portion of the mandibular arch of a 20 somite *CAG::H2B-GFP* embryo. The movie begins and ends with rostral views but the predominant view is from a distal perspective. Nuclei subtly adjusted position relative to one another but did not intercalate.

**Movie 17** Live, time-lapse, colour-coded fluorescence lifetime evaluation of vinculin tension sensor (VinTS) among epithelial cells in the mandibular arch of an intact mouse embryo.

**Movie 18** Cytosolic fluorescence intensity of the X-rhod-1 calcium indicator fluctuated asynchronously among cells in the middle region of the WT mandibular arch (see Fig. 4F, Supplementary Fig. 4G for quantification).

**Movie 19** X-rhod-1 calcium indicator fluctuation among cells in the distal region of the WT mandibular arch (see Fig. 4F, Supplementary Fig. 4G for quantification).

**Movie 20** Rendering of 3D OPT image of the right mandibular arch of a 19 somite stage *Wnt5a^-/-^* mutant.

**Movie 21** Rendering of 3D OPT image of the right mandibular arch of a 21 somite stage *Wnt5a^-/-^* mutant.

**Movie 22** Rendering of 3D OPT image of the predicted shape of the right mandibular arch of a 21 somite *Wnt5a^-/-^* mutant based on our finite element model. Four hours of growth (between 19-21 somite stages) were simulated starting with the actual shape of the 19 somite stage arch of a *Wnt5a^-/-^* mutant. Inputs included spatial distributions of cell cycle time, epithelial and mesenchymal Young’s modulus, and whole tissue viscosity.

**Movie 23** Three dimensional rendering of the right mandibular arch of a 20 somite *CAG::H2B-GFP;Wnt5a^-/-^* embryo imaged by live light sheet microscopy. Nuclei in the middle region lacked longitudinal orientation as in the WT arch (Mov. 4).

**Movie 24** A small-scale volume of several cells from within the middle region of the mandibular arch of a 20 somite *CAG::H2B–GFP;Wnt5a^-/-^* embryo. The predominant view is sagittal and distal is down and to the right. Nuclei subtly adjusted positions relative to one another but did not intercalate.

**Movie 25** Cytosolic fluorescence intensity of the Fluo-8 calcium indicator fluctuated among cells in the middle region of the WT mandibular arch (see Fig. 5E, Supplementary Fig. 5D for quantification).

**Movie 26** Fluctuation of the cytosolic fluorescence intensity of the Fluo-8 calcium indicator was dampened among cells in the middle region of the *Wnt5a^-/-^* mutant mandibular arch (see Fig. 5E, Supplementary Fig. 5D for quantification).

**Movie 27** Rendering of 3D OPT image of the right mandibular arch of a 21 somite stage transgenic *Sox9:Cre;Z/Wnt5a* embryo that ubiquitously overexpressed *Wnt5a*.

**Movie 28** Mesenchymal cell movements in a 21 somite *T:Cre;Yap^f/+^;Taz^f/f^* mutant mandibular arch. Intercellular movements are abundant but disoriented and not purely centripetal (compare with Mov. 12). Rostral (anterior) is toward to the top and distal is toward the right.

**Movie 29** Fluctuation of the cytosolic fluorescence intensity of the Fluo-8 calcium indicator was dampened among cells in the middle region of the *Piezo1^-/-^* mutant mandibular arch (see Supplementary Fig. 7A for quantification).

**Movie 30** Rendering of 3D OPT image of the right mandibular arch of a 21 somite stage *Piezo1^-/-^* mutant.

## REFERENCES

1 Etournay, R. et al. Interplay of cell dynamics and epithelial tension during morphogenesis of the Drosophila pupal wing. Elife 4, e07090, doi:10.7554/eLife.07090.

2 Bertet, C., Sulak, L. & Lecuit, T. Myosin-dependent junction remodelling controls planar cell intercalation and axis elongation. Nature 429, 667–671 (2004).

3 Blankenship, J. T., Backovic, S. T., Sanny, J. S., Weitz, O. & Zallen, J. A. Multicellular rosette formation links planar cell polarity to tissue morphogenesis. Dev Cell 11, 459–470 (2006).

4 Keller, R. E. Time-lapse Cinemicrographic AnaIysis of SuperficiaI CeII Behavior during and prior to Gastrulation in Xenopus laevis. Journal of Morphology 157, 223–248 (1978).

5 Guirao, B. et al. Unified quantitative characterization of epithelial tissue development. Elife 4, doi:10.7554/eLife.08519 (2015).

6 Irvine, K. D. & Wieschaus, E. Cell intercalation during Drosophila germband extension and its regulation by pair-rule segmentation genes. Development 120, 827–841 (1994).

7 Wallingford, J. B. et al. Dishevelled controls cell polarity during Xenopus gastrulation. Nature 405, 81–85, doi:10.1038/35011077 (2000).

8 Williams, M. L. & Solnica-Krezel, L. Regulation of gastrulation movements by emergent cell and tissue interactions. Curr Opin Cell Biol 48, 33–39, doi:10.1016/j.ceb.2017.04.006 (2017).

9 Martin, A. C., Kaschube, M. & Wieschaus, E. F. Pulsed contractions of an actin-myosin network drive apical constriction. Nature 457, 495–499, doi:10.1038/nature07522 (2009).

10 Butler, L. C. et al. Cell shape changes indicate a role for extrinsic tensile forces in Drosophila germ-band extension. Nat Cell Biol 11, 859–864, doi:10.1038/ncb1894(2009).

11 Keller, R. et al. Mechanisms of convergence and extension by cell intercalation. Philos Trans R Soc Lond B Biol Sci 355, 897–922, doi:10.1098/rstb.2000.0626 (2000).

12 Heisenberg, C. P. & Bellaiche, Y. Forces in tissue morphogenesis and patterning. Cell 153, 948–962, doi:10.1016/j.cell.2013.05.008 (2013).

13 Ninomiya, H., Elinson, R. P. & Winklbauer, R. Antero-posterior tissue polarity links mesoderm convergent extension to axial patterning. Nature 430, 364–367, doi:10.1038/nature02620 (2004).

14 Lau, K. et al. Anisotropic stress orients remodelling of mammalian limb bud ectoderm. Nat Cell Biol 17, 569–579, doi:10.1038/ncb3156 (2015).

15 Yu, J. C. & Fernandez-Gonzalez, R. Local mechanical forces promote polarized junctional assembly and axis elongation in Drosophila. Elife 5, doi:10.7554/eLife.10757.

16 Rauzi, M., Verant, P., Lecuit, T. & Lenne, P. F. Nature and anisotropy of cortical forces orienting Drosophila tissue morphogenesis. Nat Cell Biol 10, 1401–1410, doi:10.1038/ncb1798 (2008).

17 Morishita, Y., Hironaka, K. I., Lee, S. W., Jin, T. & Ohtsuka, D. Reconstructing 3D deformation dynamics for curved epithelial sheet morphogenesis from positional data of sparsely-labeled cells. Nat Commun 8, 15, doi:10.1038/s41467-017-00023-7 (2017).

18 Trichas, G. et al. Multi-cellular rosettes in the mouse visceral endoderm facilitate the ordered migration of anterior visceral endoderm cells. PLoS Biol 10, e1001256, doi:10.1371/journal.pbio.1001256 (2012).

19 Dicko, M. et al. Geometry can provide long-range mechanical guidance for embryogenesis. PLoS Comput Biol 13, e1005443, doi:10.1371/journal.pcbi.1005443 (2017).

20 Heisenberg, C. P. D’Arcy Thompson’s ‘on Growth and form’: From soap bubbles to tissue self-organization. Mech Dev 145, 32–37, doi:10.1016/j.mod.2017.03.006 (2017).

21 Moore, S. W., Keller, R. E. & Koehl, M. A. The dorsal involuting marginal zone stiffens anisotropically during its convergent extension in the gastrula of Xenopus laevis. Development 121, 3131–3140 (1995).

22 Serwane, F. et al. In vivo quantification of spatially varying mechanical properties in developing tissues. Nat Methods 14, 181–186, doi:10.1038/nmeth.4101 (2017).

23 Bi, D., Lopez, J. H., Schwarz, J. M., Manning, M. L. Energy barriers and cell migration in densely packed tissues. Soft Matter 10, 1885–1890 (2014).

24 Marmottant, P. et al. The role of fluctuations and stress on the effective viscosity of cell aggregates. Proc Natl Acad Sci U S A 106, 17271–17275, doi:10.1073/pnas.0902085106 (2009).

25 Wen, J. W. & Winklbauer, R. Ingression-type cell migration drives vegetal endoderm internalisation in the Xenopus gastrula. Elife 6, doi:10.7554/eLife.27190 (2017).

26 Shindo, A. & Wallingford, J. B. PCP and septins compartmentalize cortical actomyosin to direct collective cell movement. Science 343, 649–652, doi:10.1126/science.1243126 (2014).

27 Green, J. B. & Davidson, L. A. Convergent extension and the hexahedral cell. Nat Cell Biol 9, 1010–1015, doi:10.1038/ncb438 (2007).

28 Gao, B. et al. Wnt signaling gradients establish planar cell polarity by inducing Vangl2 phosphorylation through Ror2. Dev Cell 20, 163–176, doi:10.1016/j.devcel.2011.01.001 (2011).

29 Romereim, S. M., Conoan, N. H., Chen, B. & Dudley, A. T. A dynamic cell adhesion surface regulates tissue architecture in growth plate cartilage. Development 141, 2085–2095, doi:10.1242/dev.105452 (2014).

30 Boehm, B. et al. The role of spatially controlled cell proliferation in limb bud morphogenesis. PLoS Biol 8, e1000420, doi:10.1371/journal.pbio.1000420 (2010).

31 Mao, Q., Stinnett, H. K. & Ho, R. K. Asymmetric cell convergence-driven zebrafish fin bud initiation and pre-pattern requires Tbx5a control of a mesenchymal Fgf signal. Development 142, 4329–4339, doi:10.1242/dev.124750 (2015).

32 Wyngaarden, L. A. et al. Oriented cell motility and division underlie early limb bud morphogenesis. Development 137, 2551–2558, doi:10.1242/dev.046987 (2010).

33 Gros, J. et al. WNT5A/JNK and FGF/MAPK Pathways Regulate the Cellular Events Shaping the Vertebrate Limb Bud. Curr Biol 20 (2010).

34 Gros, J. & Tabin, C. J. Vertebrate limb bud formation is initiated by localized epithelial-to-mesenchymal transition. Science 343, 1253–1256, doi:10.1126/science.1248228 (2014).

35 Platt, J. B. Ontogenetic differentiations of the ectoderm in Necturus. Anatomischer Anzeiger [Anatomical Gazette] 9, 51–56 (1894).

36 Johnston, M. C. A radioautographic study of the migration and fate of cranial neural crest cells in the chick embryo. Anat Rec 156, 143–155, doi:10.1002/ar.1091560204 (1966).

37 Noden, D. M. An analysis of migratory behavior of avian cephalic neural crest cells. Dev Biol 42, 106–130 (1975).

38 Trainor, P. A., Tan, S. S. & Tam, P. P. Cranial paraxial mesoderm: regionalisation of cell fate and impact on craniofacial development in mouse embryos. Development 120, 2397–2408 (1994).

39 Edlund, R. K., Ohyama, T., Kantarci, H., Riley, B. B. & Groves, A. K. Foxi transcription factors promote pharyngeal arch development by regulating formation of FGF signaling centers. Dev Biol 390, 1–13, doi:10.1016/j.ydbio.2014.03.004 (2014).

40 Abe, M., Ruest, L. B. & Clouthier, D. E. Fate of cranial neural crest cells during craniofacial development in endothelin-A receptor-deficient mice. Int J Dev Biol 51, 97–105, doi:10.1387/ijdb.062237ma (2007).

41 Ota, M. S. et al. Twist is required for patterning the cranial nerves and maintaining the viability of mesodermal cells. Dev Dyn 230, 216–228, doi:10.1002/dvdy.20047 (2004).

42 Ferretti, E. et al. A conserved Pbx-Wnt-p63-Irf6 regulatory module controls face morphogenesis by promoting epithelial apoptosis. Dev Cell 21, 627–641, doi:10.1016/j.devcel.2011.08.005 (2011).

43 Akiyama, R. et al. Distinct populations within Isl1 lineages contribute to appendicular and facial skeletogenesis through the beta-catenin pathway. Dev Biol 387, 37–48, doi:10.1016/j.ydbio.2014.01.001 (2014).

44 ten Berge, D. et al. Prx1 and Prx2 are upstream regulators of sonic hedgehog and control cell proliferation during mandibular arch morphogenesis. Development 128, 2929–2938 (2001).

45 Xu, H., Cerrato, F. & Baldini, A. Timed mutation and cell-fate mapping reveal reiterated roles of Tbx1 during embryogenesis, and a crucial function during segmentation of the pharyngeal system via regulation of endoderm expansion. Development 132, 4387–4395, doi:10.1242/dev.02018 (2005).

46 Rochard, L. J., Ling, I. T., Kong, Y. & Liao, E. C. Visualization of Chondrocyte Intercalation and Directional Proliferation via Zebrabow Clonal Cell Analysis in the Embryonic Meckel’s Cartilage. J Vis Exp, e52935, doi:10.3791/52935 (2015).

47 Liu, W., Selever, J., Lu, M. F. & Martin, J. F. Genetic dissection of Pitx2 in craniofacial development uncovers new functions in branchial arch morphogenesis, late aspects of tooth morphogenesis and cell migration. Development 130, 6375–6385, doi: 10.1242/dev.00849 (2003).

48 McLennan, R., Teddy, J. M., Kasemeier-Kulesa, J. C., Romine, M. H. & Kulesa, P. M. Vascular endothelial growth factor (VEGF) regulates cranial neural crest migration in vivo. Dev Biol 339, 114–125, doi:10.1016/j.ydbio.2009.12.022 (2010).

49 Bacino, C. in GeneReviews(R) (eds R. A. Pagon et al.) (1993).

50 Person, A. D. et al. WNT5A mutations in patients with autosomal dominant Robinow syndrome. Dev Dyn 239, 327–337, doi:10.1002/dvdy.22156 (2010).

51 Roifman, M., Brunner, H., Lohr, J., Mazzeu, J. & Chitayat, D. in GeneReviews(R) (eds R. A. Pagon et al.) (1993).

52 Roifman, M. et al. De novo WNT5A-associated autosomal dominant Robinow syndrome suggests specificity of genotype and phenotype. Clin Genet 87, 34–41, doi:10.1111/cge.12401 (2015).

53 Yamaguchi, T. P., Bradley, A., McMahon, A. P. & Jones, S. A Wnt5a pathway underlies outgrowth of multiple structures in the vertebrate embryo. Development 126, 1211–1223 (1999).

54 Tuttle, A. H. et al. Immunofluorescent detection of two thymidine analogues (CldU and IdU) in primary tissue. Journal of visualized experiments: JoVE, doi:10.3791/2166 (2010).

55 Adams, D. S., Keller, R. & Koehl, M. A. The mechanics of notochord elongation, straightening and stiffening in the embryo of Xenopus laevis. Development 110, 115–130 (1990).

56 Priess, J. R. & Hirsh, D. I. Caenorhabditis elegans morphogenesis: the role of the cytoskeleton in elongation of the embryo. Dev Biol 117, 156–173 (1986).

57 Williams, M., Yen, W., Lu, X. & Sutherland, A. Distinct apical and basolateral mechanisms drive planar cell polarity-dependent convergent extension of the mouse neural plate. Dev Cell 29, 34–46, doi:10.1016/j.devcel.2014.02.007 (2014).

58 Matzke, E. B. Volume-shape relationships in variant foams. A further study of the role of surface forces in three-dimensional cell shape determination. American Journal of Botany 33, 58–80 (1946).

59 Weaire, D., Tobin, S.T., Meagher, A.J., Hutzler, S. in Foam Engineering: Fundamentals and Applications (ed P. Stevenson) Ch. 2, (John Wiley & Sons, 2012).

60 Merkel M., M. L. A geometrically controlled rigidity transition in a model for confluent 3D tissues. arXiv:1706.02656[cond-mat.soft] (2017).

61 Rosello, C., Ballet, P., Planus, E., Tracqui, P. Model driven quantification of individual and collective cell migration. Acta Biotheoretica 52, 343–363 (2004).

62 Wu, P. H., Giri, A., Sun, S. X. & Wirtz, D. Three-dimensional cell migration does not follow a random walk. Proc Natl Acad Sci U S A 111, 3949–3954, doi:10.1073/pnas.1318967111 (2014).

63 Skoglund, P., Rolo, A., Chen, X., Gumbiner, B. M. & Keller, R. Convergence and extension at gastrulation require a myosin IIB-dependent cortical actin network. Development 135, 2435–2444, doi:10.1242/dev.014704 (2008).

64 Tetley, R. J., Blanchard, G. B., Fletcher, A. G., Adams, R. J. & Sanson, B. Unipolar distributions of junctional Myosin II identify cell stripe boundaries that drive cell intercalation throughout Drosophila axis extension. Elife 5, doi:10.7554/eLife.12094 (2016).

65 Krieg, M. et al. Tensile forces govern germ-layer organization in zebrafish. Nat Cell Biol 10, 429–436, doi:10.1038/ncb1705 (2008).

66 Maitre, J. L. et al. Adhesion functions in cell sorting by mechanically coupling the cortices of adhering cells. Science 338, 253–256, doi:10.1126/science.1225399 (2012).

67 Winklbauer, R. & Parent, S. E. Forces driving cell sorting in the amphibian embryo. Mech Dev, doi:10.1016/j.mod.2016.09.003 (2016).

68 Grashoff, C. et al. Measuring mechanical tension across vinculin reveals regulation of focal adhesion dynamics. Nature 466, 263–266, doi:10.1038/nature09198 (2010).

69 Baird, M. A. et al. Local pulsatile contractions are an intrinsic property of the myosin 2A motor in the cortical cytoskeleton of adherent cells. Mol Biol Cell 28, 240–251, doi:10.1091/mbc.E16-05-0335 (2017).

70 Hunter, G. L., Crawford, J. M., Genkins, J. Z. & Kiehart, D. P. Ion channels contribute to the regulation of cell sheet forces during Drosophila dorsal closure. Development 141, 325–334, doi:10.1242/dev.097097 (2014).

71 Reginensi, A. et al. Yap– and Cdc42-dependent nephrogenesis and morphogenesis during mouse kidney development. PLoS Genet 9, e1003380, doi:10.1371/journal.pgen.1003380 (2013).

72 Reginensi, A. et al. A critical role for NF2 and the Hippo pathway in branching morphogenesis. Nat Commun 7, 12309, doi:10.1038/ncomms12309 (2016).

73 Park, H. W. et al. Alternative Wnt Signaling Activates YAP/TAZ. Cell 162, 780–794, doi:10.1016/j.cell.2015.07.013 (2015).

74 Coste, B. et al. Piezo1 and Piezo2 are essential components of distinct mechanically activated cation channels. Science 330, 55–60, doi:10.1126/science.1193270 (2010).

75 Magie, C. R., Daly, M. & Martindale, M. Q. Gastrulation in the cnidarian Nematostella vectensis occurs via invagination not ingression. Dev Biol 305, 483–497, doi:10.1016/j.ydbio.2007.02.044 (2007).

76 Kraus, Y. & Technau, U. Gastrulation in the sea anemone Nematostella vectensis occurs by invagination and immigration: an ultrastructural study. Dev Genes Evol 216, 119–132, doi:10.1007/s00427-005-0038-3 (2006).

77 Blanchard, G. B., Murugesu, S., Adams, R. J., Martinez-Arias, A. & Gorfinkiel, N. Cytoskeletal dynamics and supracellular organisation of cell shape fluctuations during dorsal closure. Development 137, 2743–2752, doi:10.1242/dev.045872 (2010).

78 David, D. J., Tishkina, A. & Harris, T. J. The PAR complex regulates pulsed actomyosin contractions during amnioserosa apical constriction in Drosophila. Development 137, 1645–1655, doi:10.1242/dev.044107 (2010).

79 He, L., Wang, X., Tang, H. L. & Montell, D. J. Tissue elongation requires oscillating contractions of a basal actomyosin network. Nat Cell Biol 12, 1133–1142, doi:10.1038/ncb2124 (2010).

80 Kim, H. Y. & Davidson, L. A. Punctuated actin contractions during convergent extension and their permissive regulation by the non-canonical Wnt-signaling pathway. J Cell Sci 124, 635–646, doi:10.1242/jcs.067579 (2011).

81 Rauzi, M., Lenne, P. F. & Lecuit, T. Planar polarized actomyosin contractile flows control epithelial junction remodelling. Nature 468, 1110–1114, doi:10.1038/nature09566 (2010).

82 Solon, J., Kaya-Copur, A., Colombelli, J. & Brunner, D. Pulsed forces timed by a ratchet-like mechanism drive directed tissue movement during dorsal closure. Cell 137, 1331–1342, doi:10.1016/j.cell.2009.03.050 (2009).

83 Maitre, J. L., Niwayama, R., Turlier, H., Nedelec, F. & Hiiragi, T. Pulsatile cell-autonomous contractility drives compaction in the mouse embryo. Nat Cell Biol 17, 849–855, doi:10.1038/ncb3185 (2015).

84 Driquez, B., Bouclet, A. & Farge, E. Mechanotransduction in mechanically coupled pulsating cells: transition to collective constriction and mesoderm invagination simulation. Phys Biol 8, 066007, doi:10.1088/1478-3975/8/6/066007 (2011).

85 Koride, S. et al. Mechanochemical regulation of oscillatory follicle cell dynamics in the developing Drosophila egg chamber. Mol Biol Cell 25, 3709–3716, doi:10.1091/mbc.E14-04-0875 (2014).

86 Wang, Q., Feng, J. J. & Pismen, L. M. A cell-level biomechanical model of Drosophila dorsal closure. Biophys J 103, 2265–2274, doi:10.1016/j.bpj.2012.09.036 (2012).

87 Machado, P. F., Blanchard, G.B., Duque, J., Gorfinkiel, N. Cytos-keletal turnover and Myosin contractility drive cell autonomous oscillations in a model of Drosophila Dorsal Closure. European Physical Journal Special Topics 223, 1391–1402 (2014).

88 Gorfinkiel, N. From actomyosin oscillations to tissue-level deformations. Dev Dyn 245, 268–275, doi:10.1002/dvdy.24363 (2016).

89 Munjal, A., Philippe, J. M., Munro, E. & Lecuit, T. A self-organized biomechanical network drives shape changes during tissue morphogenesis. Nature 524, 351–355, doi:10.1038/nature14603 (2015).

90 Julicher, F., Prost, J. Spontaneous oscillations of collective molecular motors. Physical Review Letters 78, 4510–4513 (1997).

91 Machado, P. F. et al. Emergent material properties of developing epithelial tissues. BMC Biol 13, 98, doi:10.1186/s12915-015-0200-y (2015).

92 Hadjantonakis, A. K. & Papaioannou, V. E. Dynamic in vivo imaging and cell tracking using a histone fluorescent protein fusion in mice. BMC biotechnology 4, 33 (2004).

93 Muzumdar, M. D., Tasic, B., Miyamichi, K., Li, L. & Luo, L. A global double-fluorescent Cre reporter mouse. Genesis 45, 593–605, doi:10.1002/dvg.20335 (2007).

94 Rhee, J. M. et al. In vivo imaging and differential localization of lipid-modified GFP-variant fusions in embryonic stem cells and mice. Genesis 44, 202–218, doi:10.1002/dvg.20203 (2006).

95 Ranade, S. S. et al. Piezo1, a mechanically activated ion channel, is required for vascular development in mice. Proc Natl Acad Sci U S A 111, 10347–10352, doi:10.1073/pnas.1409233111 (2014).

96 Madisen, L. et al. A toolbox of Cre-dependent optogenetic transgenic mice for light-induced activation and silencing. Nat Neurosci 15, 793–802, doi:10.1038/nn.3078 (2012).

97 Ye, X. et al. Norrin, frizzled-4, and Lrp5 signaling in endothelial cells controls a genetic program for retinal vascularization. Cell 139, 285–298, doi:10.1016/j.cell.2009.07.047 (2009).

98 Hayashi, S., Lewis, P., Pevny, L. & McMahon, A. P. Efficient gene modulation in mouse epiblast using a Sox2Cre transgenic mouse strain. Mech Dev 119 Suppl 1, S97–S101 (2002).

99 Wong, M. D., Dazai, J., Walls, J. R., Gale, N. W. & Henkelman, R. M. Design and implementation of a custom built optical projection tomography system. PLoS One 8, e73491, doi:10.1371/journal.pone.0073491 (2013).

100 Saha, R., Nix, W.D. Effects of the substrate on the determination of thin film mechanical properties by nanoindentation. Acta Mater 50, 23–38 (2002).

101 Trickey, W. R., Baaijens, F. P., Laursen, T. A., Alexopoulos, L. G. & Guilak, F. Determination of the Poisson’s ratio of the cell: recovery properties of chondrocytes after release from complete micropipette aspiration. J Biomech 39, 78–87, doi:10.1016/j.jbiomech.2004.11.006 (2006).

102 Perriot, A., Barthel, E. Elastic contact to a coated half-space: Effective elastic modulus and real penetration. Journal of Materials Research 19, 600–608 (2004).

103 Uhlenbeck, G. E., Ornstein, L.S. On the theory of the brownian motion. Physical Review 36, 823–841 (1930).

104 Wyngaarden, L. A. & Hopyan, S. Plasticity of proximal-distal cell fate in the mammalian limb bud. Dev Biol 313, 225–233, doi:10.1016/j.ydbio.2007.10.039 (2008).

105 Fernandez-Gonzalez, R. & Zallen, J. A. Oscillatory behaviors and hierarchical assembly of contractile structures in intercalating cells. Physical biology 8, 045005, doi:10.1088/1478-3975/8/4/045005 (2011).

